# Benchmarking Orthogroup Inference Accuracy: Revisiting Orthobench

**DOI:** 10.1101/2020.07.09.195586

**Authors:** D.M. Emms, S. Kelly

## Abstract

Orthobench is the standard benchmark to assess the accuracy of orthogroup inference methods. It contains 70 expert curated reference orthogroups (RefOGs) that span the Bilateria and cover a range of different challenges for orthogroup inference. Here we leveraged improvements in tree inference algorithms and computational resources to re-interrogate these RefOGs and carry out an extensive phylogenetic delineation of their composition. This phylogenetic revision altered the membership of 31 of the 70 RefOGs, with 24 subject to extensive revision and a further 7 that required minor changes. We further used these revised and updated RefOGs to provide an assessment of the orthogroup inference accuracy of widely used orthogroup inference methods. Finally, we provide an open-source benchmarking suite to support the future development and use of the Orthobench benchmark.

**Significance statement:** Orthogroup inference forms the foundation of comparative genomic analysis. Benchmarks to evaluate performance are essential to enable these methods to be compared and stimulate further method development. Here we present an update to the orthobench benchmark database and provide a comparative performance evaluation of commonly used orthogroup inference methods.

## Introduction

Determining the phylogenetic relationships between genes is fundamental to comparative biological research. While pairwise comparisons between species typically leverage the use of orthologs (i.e. genes in those species that evolved from a single ancestral gene by speciation), comparisons across multiple species require the use of orthogroups (i.e. the complete set of genes descended from a single ancestral gene in the last common ancestor of the species being analysed). Furthermore, phylogenetic analysis of orthogroups provides the basis for our understanding of the diversity and evolutionary history of life on earth. Given the fundamental utility of orthogroups in comparative biological research several automated methods have been developed to identify them from raw sequence data. Widely used methods include OrthoMCL (Li, et al. 2003), OMA (Train, et al. 2017), Hieranoid (Schreiber and Sonnhammer 2013), OrthoFinder (Emms and Kelly 2015) and SonicParanoid (Cosentino and Iwasaki 2019). Each of these methods adopt different approaches to the challenges introduced by gene duplication and loss, unequal species sampling and differential rates of sequence evolution.

Given the methodological differences between orthogroup inference methods it is important to have accurate benchmarking tools to enable their assessment. Such benchmarks are essential to enable end-users to determine which tool to use for their analysis, and to help guide developers to improve their orthogroups inference methods. The original Orthobench study (Trachana, et al. 2011) was an exemplary contribution in this endeavour. In this study the authors provided 70 expert-curated Bilateria-level orthogroups, which were termed Reference Orthogroups (RefOGs). Each of these RefOGs was intended to comprise the complete set of genes that are descended from a single copy gene in the most recent common ancestor of the Bilateria. The 70 RefOGs exemplified the range of biological and technical factors that challenge orthogroup inference methods. The authors assembled these orthogroups through expert analysis of rooted gene trees inferred from multiple sequence alignments. This benchmark database has acted as a gold standard against which orthogroup inference methods have been tested for nearly a decade.

Over the last decade tools for multiple sequence alignment and tree inference have improved considerably (Sievers and Higgins 2020). Of particular note are the improvements that have been made in phylogenetic tree inference. These include automated testing for the best fitting model of sequence evolution, new topology search strategies, higher-computational efficiency and improved parallelisation (Kalyaanamoorthy, et al. 2017; Nguyen, et al. 2015; Stamatakis 2014). These improvements, coupled with vast increases in computational power, make feasible the inference of multiple, larger gene trees using better-fitting substitution models. Unlike a decade ago, gene trees of hundreds of genes can be inferred with ease, allowing exploratory testing of phylogenetic hypotheses. This removes the need for tight inclusion thresholds which can exclude both true-positive orthogroup members as well as important context from the wider gene family necessary for accurate placement of the root of the gene tree under consideration. Thus, it is timely to re-evaluate the Orthobench benchmark database and reassess membership of its 70 RefOGs, aided by the technological advancements since the original study.

Here we utilised up-to-date bioinformatic methods to conduct an *ab initio* search, alignment and phylogenetic evaluation of the 70 RefOGs from the original Orthobench database (Trachana, et al. 2011). Using this approach, we revised the membership of 31 of the 70 RefOGs (44%). While 7 of the RefOGs required minor revision to orthogroup membership, 24 of the RefOGs required major revision that altered the phylogenetic extent of the genes in the orthogroup. To facilitate future use of this resource we provide the complete revised benchmark database and testing suite to enable orthogroup inference methods to be evaluated on this benchmark. This testing suite also includes the input proteome datasets and analysis scripts to compute the benchmark scores. Finally, to ensure reproducibility and promote further improvements to the benchmarks the complete working dataset and a summary of the evidence used to determine each updated RefOG is available from https://github.com/davidemms/Open_Orthobench

## Results

### Inference of gene trees of RefOGs in the context of their wider gene families

The first step in the assessment and potential revision of the RefOGs from Orthobench required the identification and retrieval of sets of genes from the 12 species used in the original study (Trachana, et al. 2011). Given that there had been updates to many of the genomes’ annotations since the publication of the study, the latest versions of the proteomes for the original 12 species were downloaded and are provided (Zenodo research archive https://doi.org/10.5281/zenodo.3936756 & https://github.com/davidemms/Open_Orthobench). The proteomes for an additional 3 outgroup and 2 ingroup species were also downloaded for use in this analysis. These additional species were used to provide additional evidence during the manual curation phase of the RefOG assessment but, for consistency, do not form part of the final benchmarks.

Sequence similarity searches were conducted using HMMER (Mistry, et al. 2013) with the 70 hidden Markov models prepared in the original study (Trachana, et al. 2011) used as queries. For each RefOG the set of sequences that were used for subsequent alignments and tree inference were selected directly from these HMM search results using a variable e-value threshold. Liberal e-value inclusion thresholds were used to ensure that all putative members of each orthogroup were recovered at the expense of including large numbers of false positive genes in the initial gene tree (i.e. genes that belong to the same gene family but not the target RefOG). This was done as such false positive genes are best identified and discarded on the basis of evidence provided in a gene tree, instead of on the basis of HMMER e-values. As a starting point, the highest e-value for a sequence in the RefOG tree from the original study was determined and a new e-value threshold was chosen such that there would be three times as many genes included in the gathered set than would be have been included using the e-value implied by the tree from the original study. This strategy was supported by a subsequent analysis of the final, phylogenetically-determined RefOGs. It showed that a median of 1.8 times as many genes achieved HMMER e-values better than the worst scoring true member of the RefOG as there were true members of the RefOG (Supplementary Material Figure S1).

Each gathered set of genes was subject to multiple sequence alignment using the MAFFT L-INS-I algorithm (Katoh and Standley 2013), alignment trimming using TrimAL (Capella-Gutierrez, et al. 2009) and phylogenetic inference using IQTREE (Nguyen, et al. 2015) with the best fitting model of sequence evolution inferred directly from each alignment (see Methods). This set of gene trees informed the subsequent phylogenetic analyses.

### Delineation of Bilaterian-level orthogroups from within large gene trees

The newly inferred gene trees were examined side-by-side with the trees from the original Orthobench study. In each case the Bilaterian-level orthogroup was determined and a comprehensive discussion of the evidence considered was recorded (see Supplementary Material Text S1 and https://github.com/davidemms/Open_Orthobench). Overall, 56% of RefOGs (n=39) from the previous study were confirmed with no changes to their extent. A further 10% (n=7) required only minor revision, affecting multiple genes. The remaining 34% of RefOGs (n=24) required major revision affecting the phylogenetic extent of the species with representatives in the RefOG (Fig 1A, Supplementary Material Table S1).

**Figure 1.**
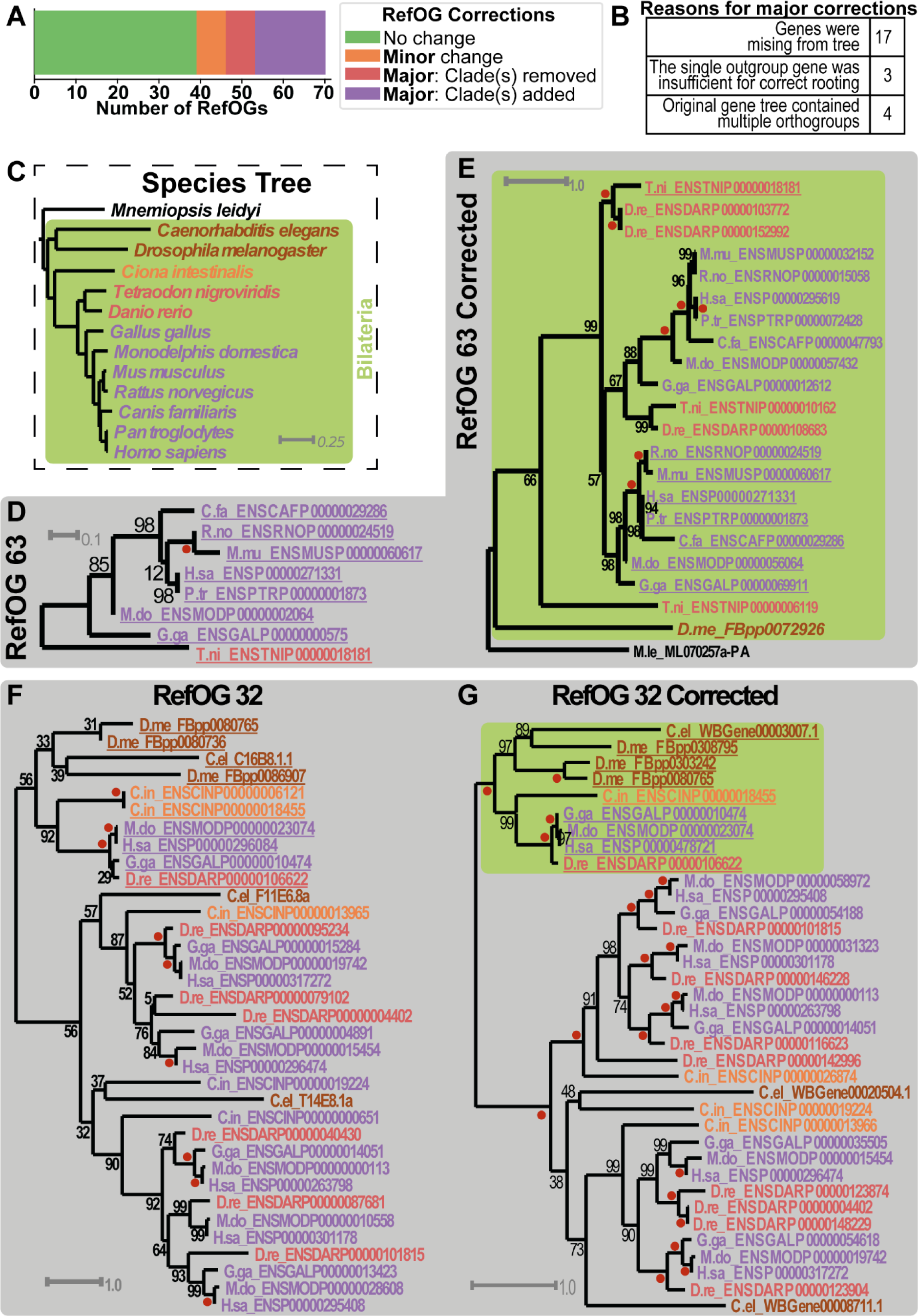
Evaluation and revision of RefOGs from Orthobench. **A)** Summary of the corrections made to the RefOG dataset. **B)** Reasons for major corrections to RefOGs from the previous study **C)** The species tree. Green shaded area shows the 12 Bilaterian species for which the Bilaterian-level orthogroups (RefOGs) were defined. One outgroup species, which appears in the gene trees in the Figure, is also shown. **D)** Example of a major improvement for which clades had been missing from the original RefOG tree: RefOG 63 gene tree as determined in the original study. **E)** Gene tree from this study showing the corrected RefOG 63 orthogroup shaded green. Phylogenetic analysis revealed a duplication within the orthogroup prior to the divergence of the vertebrates. **F)** Example of a major improvement for which extra clades of genes had been included in the original RefOG: RefOG 32 gene tree as determined in the original study. **G)** Gene tree from this study showing the corrected RefOG 32 orthogroup. Phylogenetic analysis revealed that these genes diverged from the remaining genes in the tree at a gene duplication event pre-dating the divergence of the Deuterostomes & Protostomes. Gene trees show previously identified orthogroup containing the newly delimited orthogroup from this study (green shaded clade). Genes/species are coloured according to species. Corresponding genes identified as members of the orthogroup in both studies are underlined (including when identifiers have been updated). Red dot = 100% bootstrap support.

The most common reason for major revision of a RefOG (i.e. affecting the phylogenetic extent) was that phylogenetically relevant genes were missing from the gene tree inferred in the original study. The revised phylogenetic trees support the inclusion of these genes and thus no evidence can be discerned for their original exclusion (Fig. 1B, Figs. 1C-E, Supplementary Material Table S1). It is possible that these genes did not meet the e-value threshold used for the RefOG in the original study, but the data were not available to assess this. The lenient initial threshold combined with the use of the gene trees to assess RefOG extent in the present study was designed to prevent the occurrence of such missing genes.

In addition to exclusion of clades of genes, there were also three cases of over inclusion of clades of genes (Supplementary Material Table S1). This over-inclusion was likely caused by misinterpretation of gene duplication events that occurred prior to the divergence of the Bilateria. These gene duplication events should have resulted in the identification of two Bilatarian orthogroups rather than one. This error may have occurred as there was only a single outgroup gene in the original trees and thus there was insufficient evidence to unambiguously root the tree correctly with respect to the gene duplication event (Supplementary Figure 2). In four further cases there was ambiguous or unambiguous evidence in the original tree that it contained multiple orthogroups, and further phylogenetic investigation confirmed this (Figs. 1F-G, Supplementary Material Table S1). A further 10% of RefOGs required minor revision, involving the addition or removal of single genes within already correctly identified clades. Thus, overall 44% of RefOGs required revision.

### Evaluation of orthogroup inference methods using the updated benchmarks

As the revised RefOGs were computed through manual gathering of genes and phylogenetic trees they are methodologically independent of all orthogroup inference methods. Thus, they serve as an ideal benchmark dataset on which to compare the performance of orthogroup inference methods. To facilitate the use of these RefOGs as an orthogroup benchmark a complete benchmarking suite was prepared that included the 12 input proteomes, the revised RefOGs and a script for calculating the benchmarks for a set of inferred orthogroups. A set of commonly used orthogroup inference methods comprising OrthoFinder (Emms and Kelly 2019), SonicParanoid (Cosentino and Iwasaki 2019), OrthoMCL (Li, et al. 2003), Hieranoid (Schreiber and Sonnhammer 2013) and OMA (Train, et al. 2017) were tested against the benchmarks. OrthoFinder achieved the highest F-score and recall, and obtained a more balanced precision and recall than the other methods (Fig. 2A). OMA achieved the highest precision. The ordering of the methods according to the number of RefOGs inferred exactly (no missing or extra genes) recapitulated the ordering by F-score (Fig 2B)

**Figure 2.**
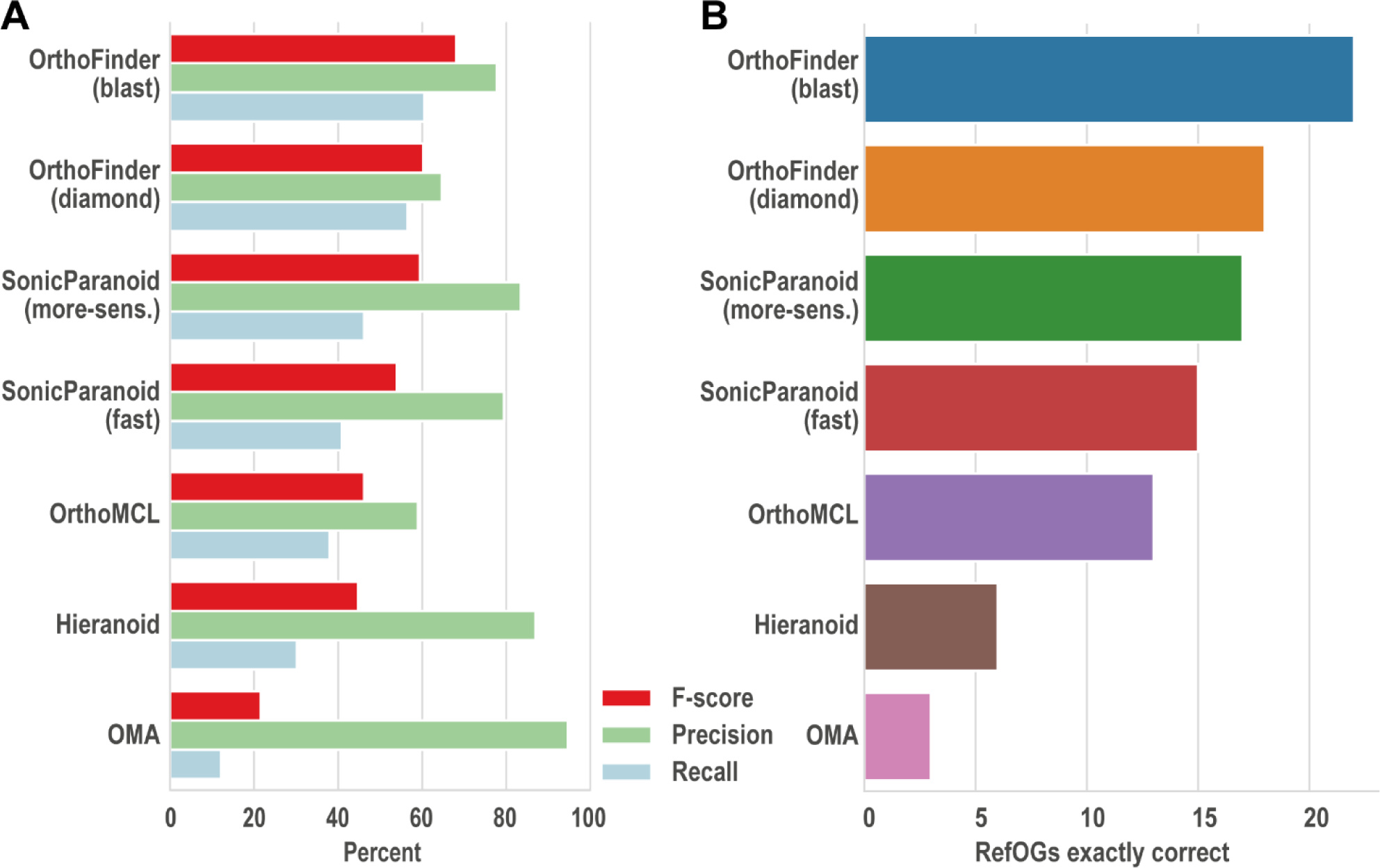
The benchmark results for the methods tested. **A)** Precision, recall and F-score. **B)** Number of orthogroups predicted exactly, with no extra or missing genes.

### An open-source benchmark to encourage future improvements

An archive of the complete set of data used to determine the members of each orthogroup is provided as supplemental data (Zenodo research archive https://doi.org/10.5281/zenodo.3936756). A GitHub repository has also been created to allow to community to rapidly identify if any further improvements should be made to the benchmarks and to allow the benchmarks to be updated accordingly: https://github.com/davidemms/Open_Orthobench. For each RefOG the following data are provided: the HMMER search results against each of the proteomes; the FASTA file of the selected sequences for subsequent tree inference; the multiple sequence alignment of these selected sequences; the rooted gene tree inferred from this alignment; and a commentary outlining the evidence considered in determining the extent of the orthogroup. The inclusion of the evidence considered allows the reasoning used in each case to be checked. It also allows any future reanalysis to take into account all the evidence already collected, and to identify what new evidence is available to support any further correction. The GitHub repository also included the input sequence files and a benchmarking script to allow any new orthogroup inference method to be tested using consistent input data and methodology.

## Discussion and Conclusions

Orthogroup inference forms the foundation of comparative genomic analysis in the post-genome era. Methods for orthogroup inference are well developed, however resources for orthogroup inference benchmarking are limited. The Orthobench (Trachana, et al. 2011) orthogroup inference benchmark has been the gold standard for benchmarking the accuracy of orthogroup inference methods for nearly a decade. In this study, we revealed that 39% of orthogroups in the Orthobench database were incorrect and needed revision. In providing these revisions we have generated a new open-source repository for the benchmarks, including all input datafiles and assessment scripts. This revised benchmark database provides more accurate assessment of orthogroup inference methods and will guide future methodological improvements.

We evaluated widely-used orthogroup inference methods on the revised benchmarks to illustrate the wide range in performance characteristics exhibited by these methods. This highlighted that, while some methods did better than others on different components of accuracy (such as precision and recall), there is still substantial room for improvement for all methods. As these methods form the foundation of thousands of comparative genomic analyses annually, such methodological improvements have the potential to drive improvements across the breath of comparative genomic research. To aid future benchmarking and accelerate any future efforts to improve the benchmarks and methods further, the complete, open-source benchmarking suite has been made available on https://github.com/davidemms/Open_Orthobench.

## Materials and methods

### Data availability and open source benchmarking repository

The input proteomes for each of the 12 species in the original Orthobench study (Trachana, et al. 2011), as well as for the three outgroup species: *Mnemiopsis leidyii, Trichoplax adhaerens* & *Nematostella vectensis* and two additional ingroup species: *Branchiostoma lanceolatum* and *Schistosoma mansoni* (diverging early within the Deuterostomes and Protostomes respectively) were downloaded from Ensembl (Cunningham, et al. 2019) in March 2020. Version numbers for all proteomes are provided in Supplemental Material Table S2. All input proteomes, HMMER search results, sequence files, alignments, trees and commentaries are provided as a supplementary data archive (Zenodo research archive https://doi.org/10.5281/zenodo.3936756). This data is also provided as an open-source GitHub repository (https://github.com/davidemms/Open_Orthobench) so that if any further improvements are identified by the community, the benchmarks can be updated accordingly. The repository also contains the script for benchmarking a set of predicted orthogroups. For each RefOG a file has been provided that details the analysis performed and the evidence that has been weighed in determining the members of each Bilaterian-level orthogroup. Users of this resource should also cite the original study (Trachana, et al. 2011).

### Reference proteomes for use in RefOG construction

As orthogroups are defined at the gene locus level, a FASTA file was prepared for each species containing the protein sequence corresponding to the longest transcript variant for each gene locus. Most orthogroup inference methods assume either implicitly or explicitly that only a single variant for each gene is included in the input files under consideration. However, both the 12 complete proteomes and the 12 ‘longest transcript variant’ proteomes are provided so that either can be used by an orthogroup inference method to test against the benchmarks. Additionally, methods that make use of outgroup species can used those employed here or any alternate ones suitable for the orthogroup inference method.

### Construction of extended gene sets for gene tree inference

The hidden Markov model (HMM) from each of the RefOGs generated in the original orthobench study was searched against the longest transcript variant proteomes using HMMER (Mistry, et al. 2013) with the command, “hmmsearch -E 0.001 –max”. As a starting point, the highest e-value for a sequence in the RefOG tree from the original study was determined and a new e-value threshold was chosen such that there would be three times as many genes included in the gathered set than would be included using the e-value implied by the tree from the original study. BLAST (Camacho, et al. 2009) and DIAMOND (Buchfink, et al. 2015) were also employed for additional searches as detailed in the evidence files associated with each RefOG. The e-value threshold, gathered set size and final RefOG size are provided in Supplementary Material Table S3.

### Inference and editing of multiple sequence alignments

Protein sequences were aligned using L-INS-I (Katoh and Standley 2013) with default parameters and trimmed using TrimAl using the options “-gt 0.5” to remove columns from the multiple sequence alignment that contained more than 50% gaps. The median percentage of gap characters for the columns removed in this way was 93% (Supplementary Material Table S3). AliView (Larsson 2014) was used to examine multiple sequence alignments. Short multiple sequence alignments (MSA) were only trimmed to remove columns with more than 75% gaps, or not at all. This requirement was judged on a case-by-case basis. The alignment lengths and number of trimmed columns for each RefOG are provided in Supplemental Table S3.

### Inference of gene trees

In all cases phylogenetic trees were inferred using IQTREE 1.6.11 using the options “-m TEST” to identify the best fitting model in each case and “-bb 1000” to perform a bootstrap analysis. The model parameters for each RefOG are provided in Supplemental Table S3. Parallelisation of multiple commands was performed using GNU Parallel (Tange 2011). Gene trees were analysed using Dendroscope (Huson and Scornavacca 2012) and the ETE library (Huerta-Cepas, et al. 2016). The complete analysis of each RefOG is detailed in Supplementary Material Text S1.

### Comparative evaluation of orthogroup inference methods

Each orthogroup inference method was run on the 12 longest transcript variant proteomes with default options. Hieranoid and OMA were additionally provided with the rooted species tree. The predicted sets of orthogroups from each of the methods were benchmarked against the RefOGs. Every pair of genes in the same predicted orthogroup that were also in the same RefOG were counted as true positives. Likewise, pairs of genes in the same predicted orthogroup but not in the same RefOG were counted as false positives (FP) and vice-versa for false negatives (FN). The precision, recall and F-score were calculated from these.

## Data Availability Statement

All data used and generated in this study are provided as a Zenodo research archive: https://doi.org/10.5281/zenodo.3936756. This same data was used to create an open-source GitHub repository (https://github.com/davidemms/Open_Orthobench) to allow any further improvements identified by the community to be incorporated into the benchmarks.

## Author Contributions

DME conceived the study and analysed the data. SK and DME wrote the manuscript.

## Acknowledgements

SK is a Royal Society University Research Fellow. This work was supported by the European Union’s Horizon 2020 research and innovation programme under grant agreement number 637765.

