## Supplementary Material for "Benchmarking Orthogroup Inference Accuracy: Revisiting Orthobench"

**Figure S1**

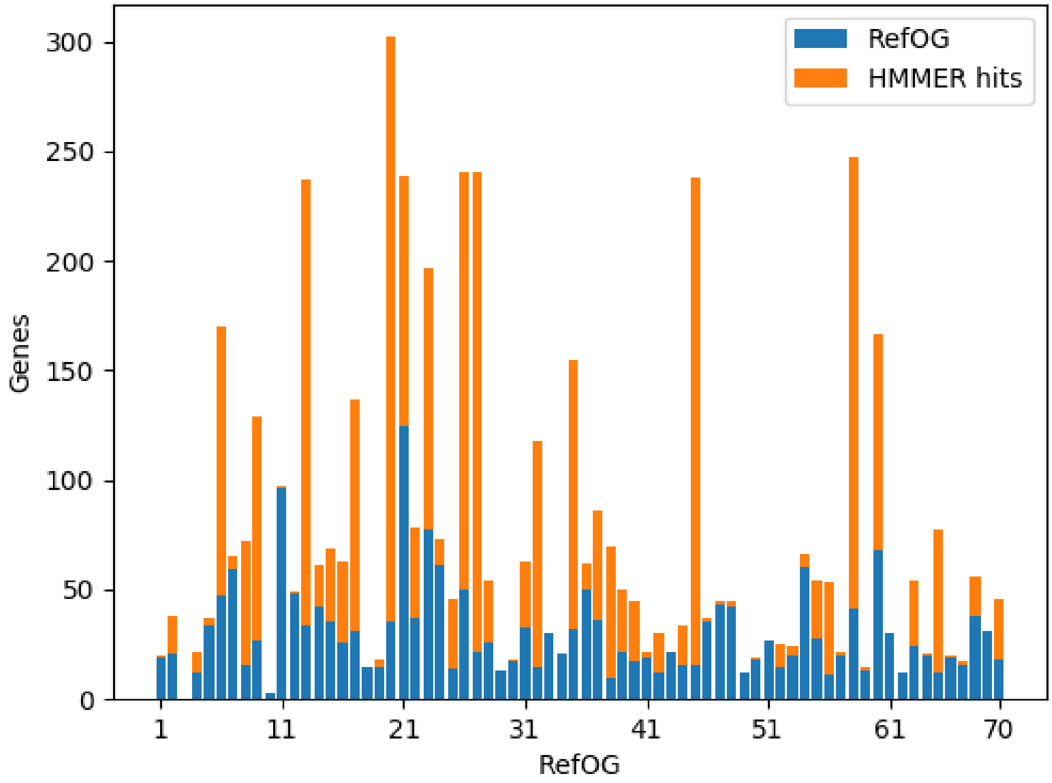


**Figure S1 Legend**

Gene trees must be inferred with a lenient HMMER e-value inclusion threshold to ensure that all actual orthogroup genes are included in the gene tree. Blue: Number of genes in each RefOG. Orange: number of genes with a HMMER hit to the RefOG profile better than or equal to the hit for the worst scoring gene that is a true member of the RefOG.

**Figure S2**

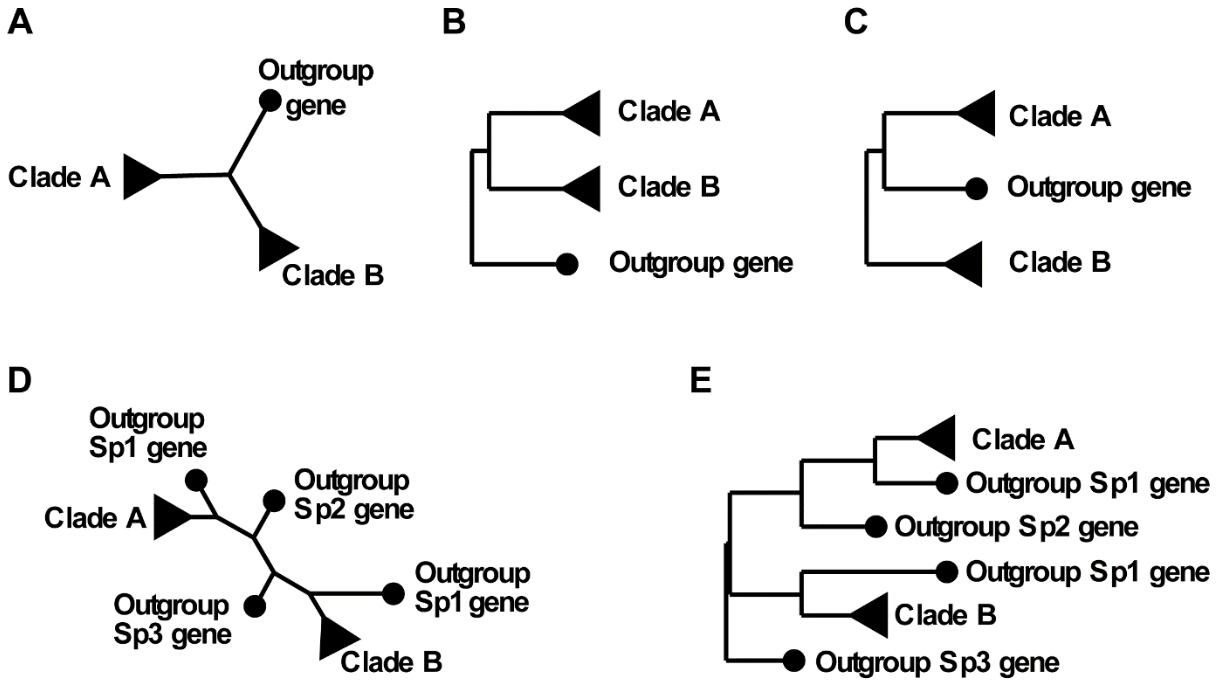


**Figure S2 Legend**

A gene tree inferred with a single outgroup gene is ambiguous if there is a duplication adjacent to the root. **A)** The inferred, unrooted gene tree showing two clades of ingroup genes from a gene duplication event either before or after the ingroup species diverged from the outgroup species. **B)** One rooting of the gene tree consistent with the species tree **C)** Another rooting of the gene tree consistent with the species tree and implying one extra gene loss event. **D)** The same hypothetical gene tree with extra outgroup genes included **E)** The gene tree can be rooted to unambiguously show the relationships between Clade A, Clade B and Outgroup Sp1 gene. Note, the original rooting ambiguity is now instead seen with respect to the relationships between Outgroup Sp3 gene and the two larger clades, but is resolved with respect to the original Clade A & Clade B.

**Table S1**

**Changes to RefOGs**

| RefOG | Change | Cause of major correction |
|---|---|---|
| 1 | Major | Genes missing from tree |
| 2 | Minor | |
| 3 | Major | Genes missing from tree |
| 4 | None | |
| 5 | None | |
| 6 | None | |
| 7 | None | |
| 8 | None | |
| 9 | Major | Genes missing from tree |
| 10 | None | |
| 11 | Major | Genes missing from tree |
| 12 | None | |
| 13 | Major | Genes missing from tree |
| 14 | Major | Genes missing from tree |
| 15 | Major | Genes missing from tree |
| 16 | None | |
| 17 | None | |
| 18 | Major | Genes missing from tree |
| 19 | None | |
| 20 | Major | Tree showed multiple orthogroups |
| 21 | Major | Genes missing from tree |
| 22 | None | |
| 23 | Minor | |
| 24 | None | |
| 25 | Minor | |
| 26 | None | |
| 27 | None | |
| 28 | None | |
| 29 | None | |
| 30 | None | |
| 31 | Major | Genes missing from tree |
| 32 | Major | Tree showed multiple orthogroups |
| 33 | None | |
| 34 | None | |
| 35 | None | |
| 36 | None | |
| 37 | Major | Single outgroup gene insufficient for correct rooting |
| 38 | None | |
| 39 | None | |
| 40 | None | |
| 41 | None | |

| | | |
|---|---|---|
| 42 | Major | Tree showed multiple orthogroups |
| 43 | None | |
| 44 | Major | Single outgroup gene insufficient for correct rooting |
| 45 | None | |
| 46 | None | |
| 47 | None | |
| 48 | Major | Genes missing from tree |
| 49 | None | |
| 50 | None | |
| 51 | Minor | |
| 52 | Major | Genes missing from tree |
| 53 | Minor | |
| 54 | Major | Genes missing from tree |
| 55 | Minor | |
| 56 | None | |
| 57 | Minor | |
| 58 | Major | Genes missing from tree |
| 59 | None | |
| 60 | Major | Tree showed multiple orthogroups |
| 61 | Major | Genes missing from tree |
| 62 | None | |
| 63 | Major | Genes missing from tree |
| 64 | None | |
| 65 | None | |
| 66 | None | |
| 67 | None | |
| 68 | Major | Single outgroup gene insufficient for correct rooting |
| 69 | Major | Genes missing from tree |
| 70 | None | |

**Table S2**

| Species | Part of benchmarks or additional | Filename | Source | Genome Assembly |
|---|---|---|---|---|
| *Caenorhabditis elegans* | In benchmark | Caenorhabditis_elegans.WBcel235.pep.all.fa | Ensembl | Wbcel235 |
| *Canis familiaris* | In benchmark | Canis_familiaris.CanFam3.1.pep.all.fa | Ensembl | 3.1 |
| *Ciona intestinalis* | In benchmark | Ciona_intestinalis.KH.pep.all.fa | Ensembl | KH |
| *Danio rerio* | In benchmark | Danio_rerio.GRCz11.pep.all.fa | Ensembl | GRCz11 |
| *Drosophila melanogaster* | In benchmark | Drosophila_melanogaster.BDGP6.28.pep.all.fa | Ensembl | BDGP6 |
| *Gallus gallus* | In benchmark | Gallus_gallus.GRCg6a.pep.all.fa | Ensembl | GRCg6a |
| *Homo sapiens* | In benchmark | Homo_sapiens.GRCh38.pep.all.fa | Ensembl | GRCh38 |
| *Monodelphis domestica* | In benchmark | Monodelphis_domestica.ASM229v1.pep.all.fa | Ensembl | ASM229v1 |
| *Mus musculus* | In benchmark | Mus_musculus.GRCm38.pep.all.fa | Ensembl | GRCm38 |
| *Pan troglodytes* | In benchmark | Pan_troglodytes.Pan_tro_3.0.pep.all.fa | Ensembl | Pan_tro_3.0 |
| *Rattus norvegicus* | In benchmark | Rattus_norvegicus.Rnor_6.0.pep.all.fa | Ensembl | Rnor_6.0 |
| *Tetraodon nigroviridis* | In benchmark | Tetraodon_nigroviridis.TETRAODON8.pep.all.fa | Ensembl | TETRAODON8 |
| *Branchiostoma lanceolatum* | Additional in-group species | Branchiostoma_lanceolatum.BraLan2.pep.all.fa | Ensembl | BraLan2 |
| *Schistosoma mansoni* | Additional in-group species | Schistosoma_mansoni.ASM23792v2.pep.all.fa | Ensembl | ASM23792v2 |
| *Nematostella vectensis* | Additional out-group species | Nematostella_vectensis.ASM20922v1.pep.all.fa | Ensembl | ASM20922v1 |
| *Trichoplax adhaerens* | Additional out-group species | Trichoplax_adhaerens.ASM15027v1.pep.all.fa | Ensembl | ASM15027v1 |
| *Mnemiopsis leidyi* | Additional out-group species | Mnemiopsis_leidyi.MneLei_Aug2011.pep.all.fa | Ensembl | MneLei_Aug2011 |

**Table S3**

| RefOG ID | Tree | HMM e-value threshold | Sequences in Gathered Field | Sequences in Final RefOG | Raw alignment length | Gap % threshold applied | Num columns removed | Ave gap % in columns removed | Final alignment length | Best fit model sequence evolution |
|---|---|---|---|---|---|---|---|---|---|---|
| 1 | RefOG001.tre | 3.30E-30 | 42 | 15 | 2734 | 50% | 2414 | 96% | 320 | LG+G4 |
| 2 | RefOG002.tre | 1.50E-26 | 60 | 19 | 16345 | 50% | 15698 | 95% | 647 | JTT+F+I+G4 |
| 3 | RefOG003.v1.tre | 8.20E-84 | 250 | 50 | 20864 | 50% | 20029 | 96% | 835 | LG+G4 |
| 3 | RefOG003.v2.tre | 1.50E-25 | 1453 | 50 | 36980 | 50% | 36641 | 98% | 339 | LG+G4 |
| 3 | RefOG003.v3.tre | 2.00E-28 | 689 | 50 | 26296 | 25% | 25993 | 97% | 303 | LG+G4 |
| 3 | RefOG003.v4.tre | 1.70E-13 | 1651 | 50 | 39812 | 25% | 39655 | 98% | 157 | LG+G4 |
| 4 | RefOG004.tre | 2.10E-87 | 39 | 12 | 2630 | 50% | 1637 | 95% | 993 | LG+G4 |
| 5 | RefOG005.tre | 1.10E-235 | 114 | 34 | 3177 | 50% | 1272 | 86% | 1905 | LG+I+G4 |
| 6 | RefOG006.tre | 1.40E-59 | 513 | 48 | 10956 | 50% | 9349 | 96% | 1607 | LG+G4 |
| 7 | RefOG007.tre | 4.10E-46 | 250 | 57 | 7852 | 50% | 7151 | 93% | 701 | WAG+F+I+G4 |
| 8 | RefOG008.tre | 3.80E-69 | 108 | 14 | 5213 | 50% | 4725 | 93% | 488 | LG+I+G4 |
| 9 | RefOG009.tre | 1.20E-66 | 153 | 23 | 13412 | 50% | 11718 | 91% | 1694 | WAG+I+G4 |
| 10 | RefOG010.tre | 6.10E-223 | 3 | 3 | 591 | 50% | 0 | 100% | 591 | JTT |
| 11 | RefOG011.tre | 7.80E-04 | 97 | 94 | 1504 | 50% | 1039 | 95% | 465 | WAG+F+I+G4 |
| 12 | RefOG012.tre | 8.20E-04 | 333 | 46 | 10087 | 50% | 9419 | 95% | 668 | LG+F+G4 |
| 13 | RefOG013.v1.tre | 6.10E-11 | 250 | 27 | 4062 | 50% | 3232 | 94% | 830 | LG+G4 |
| 14 | RefOG014.v1.tre | 9.00E-03 | 78 | 44 | 8927 | 50% | 8315 | 90% | 612 | JTT+F+G4 |
| 14 | RefOG014.v2.tre | 2.70E-04 | 39 | 44 | 3675 | 50% | 2724 | 79% | 951 | JTT+F+G4 |
| 15 | RefOG015.tre | 3.60E-05 | 73 | 30 | 6704 | 50% | 5543 | 87% | 1161 | LG+I+G4 |
| 16 | RefOG016.tre | 1.70E-43 | 75 | 25 | 3271 | 50% | 2697 | 89% | 574 | JTT+F+I+G4 |
| 17 | RefOG017.tre | 2.30E-19 | 180 | 29 | 6707 | 50% | 6068 | 90% | 639 | LG+G4 |
| 18 | RefOG018.tre | 0.0006 | 15 | 12 | 165 | 50% | 95 | 88% | 70 | VT+G4 |
| 19 | RefOG019.tre | 4.40E-08 | 20 | 13 | 3301 | 50% | 3191 | 92% | 110 | JTTDCMut+G4 |
| 20 | RefOG020.v1.tre | 8.20E-71 | 340 | 28 | 8191 | 50% | 6921 | 95% | 1270 | VT+I+G4 |

| | | | | | | | | | | |
|---|---|---|---|---|---|---|---|---|---|---|
| 20 | RefOG020.v2.tre | 1.80E-72 | 95 | 28 | 5144 | None | 0 | N/A | 5144 | VT+I+G4 |
| 21 | RefOG021.tre | 1.60E-09 | 250 | 125 | 17980 | 50% | 17479 | 93% | 501 | JTT+F+G4 |
| 22 | RefOG022.v1.tre | 0.00075 | 107 | 35 | 2826 | 50% | 2431 | 92% | 395 | DCMut+F+G4 |
| 22 | RefOG022.v2.tre | 2.20E-11 | 42 | 35 | 2196 | 50% | 1742 | 95% | 454 | Dayhoff+F+I+G4 |
| 23 | RefOG023.v1.tre | 0.00099 | 458 | 69 | 36480 | 50% | 35884 | 97% | 596 | WAG+G4 |
| 23 | RefOG023.v2.tre | 0.00099 | 170 | 69 | 19458 | 50% | 18365 | 94% | 1093 | WAG+I+G4 |
| 23 | RefOG023.v3.tre | 1.30E-09 | 101 | 69 | 14042 | 25% | 13046 | 87% | 996 | FLU+F+I+G4 |
| 24 | RefOG024.tre | 1.80E-32 | 103 | 51 | 7209 | 50% | 6800 | 97% | 409 | WAG+I+G4 |
| 25 | RefOG025.tre | 7.90E-46 | 90 | 13 | 1045 | 50% | 552 | 90% | 493 | LG+I+G4 |
| 26 | RefOG026.tre | 2.20E-88 | 250 | 43 | 53681 | 50% | 52725 | 96% | 956 | WAG+F+G4 |
| 27 | RefOG027.tre | 1.50E-64 | 250 | 20 | 42655 | 50% | 42206 | 96% | 449 | LG+F+I+G4 |
| 28 | RefOG028.tre | 1.70E-04 | 96 | 24 | 3259 | 50% | 2715 | 91% | 544 | JTT+F+G4 |
| 29 | RefOG029.v1.tre | 0.0034 | 26 | 13 | 1700 | 50% | 1375 | 92% | 325 | VT+G4 |
| 29 | RefOG029.v2.tre | 2.60E-37 | 17 | 13 | 528 | None | 0 | N/A | 528 | JTT+I+G4 |
| 30 | RefOG030.tre | 4.90E-04 | 46 | 13 | 382 | 50% | 239 | 94% | 143 | rtREV+G4 |
| 31 | RefOG031.tre | 2.20E-48 | 99 | 32 | 2175 | 50% | 1501 | 90% | 674 | JTT+F+I+G4 |
| 32 | RefOG032.v1.tre | 1.80E-86 | 250 | 14 | 6351 | 50% | 5643 | 92% | 708 | LG+F+I+G4 |
| 32 | RefOG032.v2.tre | 1.80E-86 | 105 | 14 | 2836 | 50% | 2050 | 84% | 786 | LG+F+I+G4 |
| 33 | RefOG033.v1.tre | 5.50E-90 | 87 | 30 | 1167 | 50% | 662 | 96% | 505 | LG+I+G4 |
| 33 | RefOG033.v2.tre | 2.40E-44 | 250 | 30 | 1808 | 50% | 1333 | 96% | 475 | LG+I+G4 |
| 34 | RefOG034.tre | 1.70E-19 | 66 | 21 | 648 | 50% | 375 | 91% | 273 | LG+I+G4 |
| 35 | RefOG035.tre | 1.70E-42 | 198 | 32 | 8471 | 50% | 6889 | 94% | 1582 | LG+F+I+G4 |
| 36 | RefOG036.tre | 4.20E-21 | 66 | 46 | 761 | 50% | 353 | 94% | 408 | LG+I+G4 |
| 37 | RefOG037.v1.tre | 0.00095 | 231 | 34 | 29979 | 25% | 29822 | 97% | 157 | VT+G4 |
| 37 | RefOG037.v2.tre | 2.30E-05 | 68 | 34 | 3534 | 50% | 2601 | 93% | 933 | VT+I+G4 |
| 38 | RefOG038.tre | 1.60E-42 | 123 | 9 | 13443 | 25% | 12851 | 86% | 592 | WAG+I+G4 |
| 39 | RefOG039.tre | 3.30E-89 | 249 | 20 | 60449 | 50% | 59181 | 95% | 1268 | VT+G4 |
| 40 | RefOG040.tre | 2.90E-17 | 132 | 13 | 5633 | 50% | 4539 | 86% | 1094 | LG+I+G4 |
| 41 | RefOG041.tre | 1.50E-53 | 22 | 17 | 247 | 50% | 69 | 91% | 178 | VT+G4 |
| 42 | RefOG042.v1.tre | 0.00089 | 43 | 11 | 555 | 50% | 420 | 90% | 135 | WAG+G4 |

| | | | | | | | | | | |
|---|---|---|---|---|---|---|---|---|---|---|
| 42 | RefOG042.v2.tre | 3.90E-10 | 41 | 11 | 555 | 50% | 417 | 90% | 138 | WAG+I+G4 |
| 42 | RefOG042.v3.tre | 4.70E-20 | 24 | 11 | 436 | 50% | 277 | 86% | 159 | WAG+G4 |
| 43 | RefOG043.tre | 2.20E-05 | 80 | 17 | 1827 | 50% | 1543 | 88% | 284 | WAG+G4 |
| 44 | RefOG044.tre | 2.50E-05 | 34 | 14 | 1578 | 50% | 1042 | 93% | 536 | LG+I+G4 |
| 45 | RefOG045.v1.tre | 5.20E-07 | 732 | 13 | 4422 | 25% | 4196 | 98% | 226 | LG+G4 |
| 45 | RefOG045.v2.tre | 2.30E-13 | 79 | 13 | 1383 | None | 0 | N/A | 1383 | LG+I+G4 |
| 46 | RefOG046.tre | 4.00E-17 | 37 | 32 | 154 | 50% | 62 | 94% | 92 | JTT+G4 |
| 47 | RefOG047.tre | 1.20E-12 | 45 | 36 | 750 | 50% | 358 | 96% | 392 | LG+F+G4 |
| 48 | RefOG048.tre | 2.50E-12 | 45 | 40 | 1461 | 50% | 1377 | 95% | 84 | JTT+G4 |
| 49 | RefOG049.tre | 5.70E-05 | 20 | 10 | 678 | 50% | 138 | 88% | 540 | LG+G4 |
| 50 | RefOG050.tre | 2.60E-24 | 19 | 14 | 632 | 50% | 168 | 94% | 464 | LG+G4 |
| 51 | RefOG051.tre | 0.00086 | 69 | 17 | 1154 | 50% | 953 | 91% | 201 | LG+G4 |
| 52 | RefOG052.tre | 3.00E-43 | 48 | 15 | 3295 | 50% | 2234 | 86% | 1061 | JTT+I+G4 |
| 53 | RefOG053.v1.tre | 8.70E-07 | 24 | 13 | 2982 | None | 0 | N/A | 2982 | LG+F+G4 |
| 53 | RefOG053.v2.tre | 8.60E-02 | 46 | 13 | 4776 | None | 0 | N/A | 4776 | WAG+F+G4 |
| 54 | RefOG054.tre | 0.0068 | 66 | 44 | 1762 | 50% | 1250 | 93% | 512 | VT+G4 |
| 55 | RefOG055.v1.tre | 9.60E-03 | 60 | 29 | 1473 | 50% | 1306 | 93% | 167 | JTT+G4 |
| 55 | RefOG055.v2.tre | 5.40E-04 | 52 | 29 | 1076 | 25% | 949 | 91% | 127 | JTT+F+G4 |
| 56 | RefOG056.tre | 0.00088 | 11 | 9 | 6675 | 50% | 5819 | 89% | 856 | JTT+F+I+G4 |
| 57 | RefOG057.tre | 9.50E-03 | 22 | 16 | 478 | 50% | 224 | 88% | 254 | LG+I+G4 |
| 58 | RefOG058.v1.tre | 7.40E-06 | 250 | 50 | 5172 | 50% | 4863 | 96% | 309 | VT+G4 |
| 58 | RefOG058.v2.tre | 9.80E-04 | 363 | 50 | 9572 | 50% | 9283 | 96% | 289 | WAG+G4 |
| 58 | RefOG058.v3.tre | 9.70E-04 | 93 | 50 | 2953 | 50% | 2619 | 94% | 334 | VT+I+G4 |
| 59 | RefOG059.tre | 0.0066 | 15 | 10 | 1785 | 50% | 485 | 91% | 1300 | JTT+F+G4 |
| 60 | RefOG060.v1.tre | 0.00098 | 425 | 68 | 2650 | 50% | 2464 | 97% | 186 | LG+F+G4 |
| 60 | RefOG060.v2.tre | 3.10E-06 | 250 | 68 | 1816 | 50% | 1615 | 95% | 201 | JTT+F+G4 |
| 61 | RefOG061.tre | 0.0095 | 67 | 30 | 5220 | 50% | 4625 | 90% | 595 | mtInv+F+I+G4 |
| 62 | RefOG062.tre | 9.80E-03 | 40 | 11 | 1188 | 50% | 766 | 84% | 422 | LG+G4 |
| 63 | RefOG063.v1.tre | 0.00085 | 54 | 21 | 2318 | None | 0 | N/A | 2318 | PMB+G4 |
| 63 | RefOG063.v2.tre | 9.60E-02 | 67 | 21 | 2216 | None | 0 | N/A | 2216 | PMB+I+G4 |

| | | | | | | | | | | |
|---|---|---|---|---|---|---|---|---|---|---|
| 63 | RefOG063.v3.tre | 7.30E-01 | 87 | 21 | 1897 | None | 0 | N/A | 1897 | JTT+G4 |
| 64 | RefOG064.tre | 6.80E-03 | 21 | 15 | 1479 | 50% | 1064 | 90% | 415 | JTT+F+G4 |
| 65 | RefOG065.v1.tre | 9.00E-16 | 77 | 10 | 4346 | 25% | 4133 | 83% | 213 | JTT+F+I+G4 |
| 65 | RefOG065.v2.tre | 9.00E-16 | 17 | 10 | 4346 | 25% | 3921 | 62% | 425 | JTT+F+I+G4 |
| 66 | RefOG066.tre | 1.40E-10 | 20 | 14 | 503 | 50% | 227 | 92% | 276 | LG+G4 |
| 67 | RefOG067.tre | 4.60E-09 | 18 | 13 | 347 | 50% | 222 | 93% | 125 | LG+G4 |
| 68 | RefOG068.tre | 0.0046 | 56 | 32 | 5475 | 50% | 3810 | 90% | 1665 | JTT+F+I+G4 |
| 69 | RefOG069.tre | 0.0042 | 31 | 31 | 476 | 50% | 320 | 92% | 156 | JTT+F+I+G4 |
| 70 | RefOG070.tre | 5.70E-06 | 154 | 13 | 3903 | 50% | 3489 | 92% | 414 | LG+G4 |

**Text S1: Evidence Considered**

**RefOG001.txt**

The RefOG tree is unrooted, it has been rooted on the Ciona intestinalis gene. It doesn't contain any outgroup genes to aid the analysis of the orthogroup.

The newly inferred tree shows 3 metazoan-level orthogroups, it has been rooted on the root of the one containing the target bilaterian orthogroup. It shows that the bilaterian orthogroup extends to the Protostomes, with representatives from each of the species used in this study. The order of branching at the root of the orthogroup does not exactly match expectations, with a Nematostella gene as a sister to Branchiostoma lanceolatum and in a clade with the Ciona intestinalis gene. Additionally, the three outgroup species genes are shown as more closely related to the Deuterostome clade that the Protostome genes are. This is not strong evidence of the Protostome genes being part of a separate orthogroup for a number of reasons.

1. The bootstrap support values supporting the topology are not high 2. There is no evidence of a gene duplication giving rise to this additional clade 3. The clade has likely been incorrectly rooted by the orthogroups used as an outgroup. These outgroups are separated by a particularly long branch (this can be best seen with a Radial Phylogram view). and the intersection of this long branch with the very short branches at the root of the clade of interest is highly likely to be inaccurate. Extracting just this sub-tree and rooting it on the gene from the Ctenophore Mnemiopsis leidyi resolves this confusion to come extent (v2 tree).

**RefOG002.txt**

The RefOG tree is unrooted, it has been rooted on the C. elegans gene, which is the earliest diverging gene in the tree.

The newly inferred tree contains a single outgroup Mnemiopsis leidyi gene, on which it has been rooted. This tree appears to show three bilaterian orthogroups. The target orthogroup and its sister orthogroup both contain genes from the Deuterostomes and from C. elegans. The tree is in good agreement with the orthogroup from the previous study, except that the original study missed a Drosophila gene or it was unannotated in its input data.

Although both the C. elegans and Drosophila genes are separated from the remainder of the orthogroup by a Trichoplax adhaerens gene, this is not good evidence for a duplication prior to the origin of the orthogroup and the C. elegans & Drosophila genes being part of a separate orthogroup. This would require a gene duplication event and subset loss of the Deuterostome clade from this hypothetical orthogroup in addition to loss of the Protostome clade from the target orthogroup. The clade with Trichoplax has only 79% bootstrap support, and these short, early branches in metazoan gene trees are often not resolved correctly as can be seen in many of the gene trees presented here. Similarly, the topology of the corresponding species tree has been challenging to reconstruct even when using molecular data from many gene families.

**RefOG003.txt**

RefOG is rooted on a gene from Hydra magnipapillata.

The RefOG tree is troubling since the poorest hit of the hmmer profile for any gene in the tree was 1.8e-25 whereas there are 1453 genes in the new study which are as good a match to the profile as this. The RefOG tree only contains 99 genes.

1st newly inferred tree

===================

A first attempt at a new tree for the orthogroup contained 250 genes. It shows that there are two clades from the original orthogroup that are separated by a large distance and a large number of other clades within the larger gene family. This shows clearly a number of bilaterian orthogroups that are homologous to the target orthogroup, but the resolution of the status of the target orthogroup is not entirely clear. To ensure that there are no false positives (genes missing from the tree that should be included) a new tree has been inferred with all 1453 genes with as good a hit as the poorest match to the hmm profile from the original tree. This should ensure that all relevant clades are captured in their entirety and so will allow for a clearer interpretation of the target orthogroup. If the orthogroup is of a still larger extent then a larger tree can be inferred.

Second tree

==========

Clade A

----------

The second tree contains the genes from the wider gene family. It shows that the suggested RefOG from the previous study come from two very separate orthogroups and a few genes from elsewhere within the gene family. The true bilaterian-orthogroup containing Homo_sapiens_ENSP00000419199 (Clade A) appears to be the best fit to the profile used to search for members of the orthogroup. Approximately 50% of its members are genes unidentified in the previous study. It is well-defined by the new gene tree and contains four clades originating from duplications in the MRCA of the vertebrates and stretches back to representatives in C. elegans. It has 84% bootstrap support in the FastTree tree.

Clade B

----------

The tree also shows the second clade that contains genes identified in the previous study, e.g. Homo_sapiens_ENSP00000350331. It is made up of two clades which originated at a duplication at the base of the Deuterostomes. It appears to have had at least one or two duplications at the same location. The presence of C. elegans and Drosophila genes in the orthogroup is not clear from an initial FastTree tree, and the full maximum likelihood tree will be required.

The RefOG was built around COG0666, Ankyrin repeat and SOCS box-containing (ASB) family of proteins. This appears to be defined based on a family of genes within the Fungi, if so it may be that neither of the two clades from the original study is closer to the COG than the other, or one may be more closely related to COG0666 than the other.

A number of analyses were performed that were not ultimately successful.

Reanalysis

=========

A reanalysis of this RefOG was performed. A new hmm profile was built around 48 genes from the clade identified as clearly part of the orthogroup in the previous analyses (Clade A, with human genes: ENSP00000499751, ENSP00000397655, ENSP00000419199, ENSP00000451575 & ENSP00000480388). There were a very large number of good hits. An e-value cut-off of 10^-40 was chosen for a tentative first tree to see if it appeared likely to contain all genes which could potentially belong to the orthogroup. This produced 689 hits. These were aligned with mafft linsi, trimmed of columns with greater than 75% gaps and a tree was inferred using iqtree. The tree (v3) was rooted on

metazoan-level orthogroup that was distant in the tree from the genes of interest. The target RefOG appeared relatively clear in this tree. There were a number of duplications near the base of the Deuterostomes and an outgroup gene (Mnemiopsis_leidyi_ML358816a-PA) at the base of the bilaterian orthogroup with 99% bootstrap support containing approximately 52 genes.

The pattern of e-values for the hits in the tree was initially unexpected. All genes in the bilaterian orthogroup had e-values for their hits mostly between 1e-170 and 1e-260. The hit for the outgroup gene was 1e-63 and for the sister orthogroup most hits were around 1e-45. So far this is as would be expected. However, there were three further clades throughout the tree (but at some distance) with e-values around 1e-170. This was explained by examination of the MSA. Genes with poorer hits had lost most of the sequence after approximately column 590 in the trimmed MSA, those with better hits had not. This explained the apparently distant clades that had been included in the RefOG from the original study to the exclusion of many genes more closely related in the tree. These genes had scored good hits whereas the more closely related genes had been missed in the search phase because of the lower hits due to part of the gene sequence having been lost. The loss of this part of the gene sequence appears to result in an e-value of around 1e-50.

To ensure no true members of the bilaterian orthogroup had been lost due to the e-value cut-off a second tree was inferred with a cut-off of 1e-20, producing 1651 hits. It was aligned with mafft and trimmed of columns with greater than 75% gaps and a tree was inferred using IQTREE. In this tree (v4) no new genes were placed in the clade of interest in this tree. This RefOG is, however, the least certain of the 70. In the new tree there is an extra clade that could arguably be part of the RefOG according to the tree (containing ENSP00000468354). However, this clade was placed elsewhere in the v2 tree. It is hard to have confidence in the placement of the various clades in the tree of 1651 genes and so the RefOG according to v2 will be used. The v3 tree was, however, useful in determining that no genes had been missed from *within* these clades. The exact membership of this orthogroup has only been established with low confidence.

**RefOG004.txt**

The RefOG tree is rooted on a Hydra gene and shows a bilaterian orthogroup with mostly single-copy genes.

The newly inferred tree contains a single gene from the outgroup species and has been rooted on that. It shows two bilaterian level orthogroups. The branch leading to the target orthogroup is comparatively long, but both these orthogroups appear to be individually rooted correctly and the tree appears to be reliable for the analysis of the target orthogroup. It gives good agreement with the orthogroup identified in the original study.

**RefOG005.txt**

The RefOG tree is unrooted, it has rooted on the c. elegans gene.

The newly inferred tree shows the target orthogroup as part of a larger gene family and has been rooted at the root of the metazoan orthogroup that contains it. It provides strong support for the orthogroup identified in the previous study. Although the tree shows the C. elegans & Drosophila genes diverging before the Trichoplax adhaerens, Nematostella vectensis & Mnemiopsis leidyi genes this is because of the long branch from the relatively distant sister metazoan clades (from ancient duplications). It has likely intercepted the short branches as the base of the target orthogroup inaccurately. This can be best seen using (e.g. in Dendroscope) the Radial Phylogram view. Extracting just the sub-clade and rooting on the Mnemiopsis gene shows a gene tree with topology matching expectations exactly and with high bootstrap support throughout (see final tree).

**RefOG006.txt**

The RefOG tree is rooted on a Hydra gene, it has no branch lengths.

The newly inferred tree shows the target orthogroup as part of a larger gene family and has been rooted at the root of the metazoan orthogroup that contains it. It provides good support for the orthogroup identified in the original study.

**RefOG007.txt**

The RefOG tree is rooted on a Hydra magnipapillata gene and shows an orthogroup containing a duplication early in the Deuterostomes.

The newly inferred tree shows the target orthogroup as part of a larger gene family and has been rooted at the base of the metazoan orthogroup containing it. It provides good support for the orthogroup defined in the previous study. The target orthogroup and its sister orthogroup are both demarcated by genes from the Cnidarian Nematostella, and the target orthogroup has 100% bootstrap support.

**RefOG008.txt**

The RefOG tree is rooted on a Nematostella gene and shows an orthogroup of single-copy genes present in all species.

The newly inferred tree shows a clear metazoan orthogroup as part of a larger gene family, it has been rooted at the root of this metazoan orthogroup. It provides strong support for the orthogroup from the original study. As with the previous study the Ciona gene has been judged to be part of the orthogroup despite being placed within a clade with Trichoplax and Nematostella with low bootstrap support.

**RefOG009.txt**

The RefOG tree is rooted on Hydra_magnipapillata_Hma2.217448 and shows a single orthogroup containing a duplication at the base of the vertebrates.

The newly inferred tree shows the wider gene family and has been rooted at the base of the Metazoan orthogroup that contains the target bilaterian orthogroup and a sister orthogroup. It is in good agreement with the tree from the previous study, but shows additionally the Drosophila and C. elegans that also belong to the orthogroup. This is well supported, the target orthogroup has 100% bootstrap support, as does the sister orthogroup, which also contains its C. elegans & Drosophila genes.

**RefOG010.txt**

There are 3 sequences within the input dataset with significant hits (e < 0.001). These 3 sequences have a maximum e-value of 1e-223. These are the mouse and rat genes from the original study. The remaining two genes identified in the original study have had their models retired, although the evolutionary conservation identified in the original study suggests that there is a good possibility that there are true genes. The new RefOG, for the purpose of benchmarking orthogroup inference methods based on the companion input proteome files will include just the three genes existing in these proteomes.

**RefOG011.txt**

The RefOG tree is unrooted, it consists of a number of clades with the earliest diverging species being Gallus gallus. It has been rooted on one of these clades, although it's not possible to know which branch represents the oldest duplication.

The newly inferred tree recovers the whole metazoan-orthogroup back to Mnemiopsis leidyi. It provides a plausible rooting of the RefOG from the previous study and shows to earlier duplications also within the orthogroup, both in the common ancestor of the vertebrates. It also recovers two genes in C. elegans and one in Ciona intestinalis that are also likely members of the orthogroup. There is some evidence against the inclusion of these genes in that there is a Nematostella vectensis gene shown as more closely related to the clade of genes than they are. However, the bootstrap support for this is 75%, and there is no evidence for a pre-bilaterian duplication giving rise to just these three genes as a separate orthogroup. They have therefore also been included, with moderate confidence.

**RefOG012.txt**

The RefOG tree has been rooted on a Nematostella gene. It shows a relatively straightforward orthogroup with presence in all or most species.

The tree shows multiple metazoan level orthogroups, it has been rooted at the base of a pair of metazoan orthogroups, with one of them containing the target RefOG. It shows good agreement with the orthogroup from the original study with 100% bootstrap support, with updating of genes to the new gene models.

**RefOG013.txt**

The original RefOG consists entirely of genes from Drosophila and so there was no extra information that could be used to root it.

It was found that progressively wider reaching trees were required to correctly interpret this orthogroup. In these trees it was eventually found that the orthogroup was of quite limited extent, as found in the original study, although it was found in this new study that there were also genes from C. elegans in the orthogroup:

Inferred Tree v1:

The tree shows clearly that these genes are part of a large gene family that has duplicated frequently in other species in addition to the Duplications identified in Drosophila in the original study. The correct rooting of the tree is non-trivial since it contains a number of metazoan-orthogroups each with a selection of genes from the outgroup species used in this study. The branch lengths give quite a clear delineation of a number of these orthogroups (view the tree as a radial phylogram and highlight all the outgroup genes to see this). The target orthogroup is the most divergent within the gene family and the most challenging to analyse. The tree has been rooted on a well-defined metazoan level orthogroup. This orthogroup has genes from Trichoplax adhaerens, Nematostella vectensis and the ingroup species. The inspection of the tree shows that rooting on any of the clear metazoan-orthogroups should allow the target clade to be analysed correctly (since the root will be correctly placed with respect to all the genes of interest). The rooting will be re-evaluated again at the end of the analysis to ensure that other possible rootings have no effect on the interpretation of the target orthogroup.

The tree shows clearly a metazoan-orthogroup that encompasses the target bilaterian-orthogroup (the clade containing 131 genes & with 87% bootstrap support). It contains early-diverging clades of genes from Mnemiopsis leidyi & Trichoplax adhaerens. It is not clear if the branch from the other orthogroups correct roots this Metazoan-orthogroup since an alternative rooting (version_2) is more parsimonious. Nevertheless, both this tree and the version_2 tree show strong evidence that there was a gene duplication event prior to the origin of the orthogroup that gave rise to the target orthogroup (only in the Protostomes & Branchiostoma) and a sister orthogroup (in the Protostomes and Deuterostomes). The bootstrap support is high for this and there are many genes confirming this (from C. elegans, Drosophila, Branchiostoma & Schistosoma). This confirms part of the original study, in excluding and vertebrate species from the orthogroup, but identified that there are many more C. elegans and Drosophila genes that are also part of the orthogroup. The question of rooting, raised above, does not alter this interpretation.

**RefOG014.txt**

RefOG tree is unrooted. It has been rooted on the branch separating the Danio and Tetraodon genes from the remaining vertebrate genes. There are no earlier branches.

The tree for the default search appeared incomplete and so a wider search with a less stringent e-value cut-off was used.

Inferred tree v1:

The tree contains a clear clade, more ancient than the target orthogroup and the tree additionally contains some more distantly related or false positive genes. A new tree will be inferred on the clade of interest. The tree shows a sub-clade of 20 Human and Chimp genes containing Homo_sapiens_ENSP00000477979 which will be excluded from the new tree as spurious hits. They have less good hits to the hmm profile, but this decision is with low certainty.

Inferred tree 2:

The new tree has been inferred and rooted on Nematostella_vectensis_EDO31099, the earliest diverging gene in the clade. The new tree shows a number of new genes within the tetrapods that are clearly members of the orthogroup and were either missed in the earlier study or are new gene models. The tree shows two clades that duplicated before the root of the orthogroup. The placement of the duplication before the orthogroup rather than within it is with low confidence since the only evidence is the positioning of a single Drosophila gene within one of these post-duplication clades but with only 24% bootstrap support. This is also the most conservative interpretation since it agrees with the previous study. No genes have changed position with respect to the orthogroup between the two trees.

Revisiting the clade of 20 Pan & Homo genes: Re-examining the original tree, there is no good reason to exclude these genes. Their placement suggests that they are a divergent clade that originated from within the orthogroup. The alignment clearly has enough phylogenetically informative columns with which to place these genes and they are also clearly homologous to the target orthogroup (also confirmed by profile search: e-value of ~ 1e-46). As before, this is not with certainty. As a result, a file is included listing both genes that are in the orthogroup but with low certainty (Homo/Pan clade) and genes which have been excluded from the orthogroup with low certainty.

**RefOG015.txt**

The RefOG shows an orthogroup with genes in Danio rerio, Gallus gallus & Tetraodon nigroviridis. It has a duplication in the common ancestor of these species, giving rise to two clades. It has been rooted on this gene duplication event.

The newly inferred tree shows the orthogroup to be part of a larger gene family. There are many extra Danio genes that are clearly part of the orthogroup, they may be new gene models (inspecting the Danio gene models used in the original study shows this not to be the case, at least for a number of the genes). The tree has been rooted on an ancient duplication separating two metazoan-orthogroups. The target orthogroup is part of one of these metazoan orthogroups.

There are genes from C. elegans & Drosophila which may or may not be members of the target orthogroup. There was:

Either a duplication before the origin of the orthogroup and then:

1. In one branch: a loss of Deuterostome clade after the divergence of Branchiostoma.

2. In the other branch a loss of the Protostome clade and a loss in Branchiostoma

Also, losses in Nematostella and Trichoplax in one clade. Or, more likely, the topology is not correct at this point.

Alternatively, there was no duplication, the topology is not correct at this point and the Protostome clade and Branchiostoma genes belong in the orthogroup.

The species sets match perfectly for this second alternative and it is a very tidy explanation. Arguing against this, there is a clade with 100% bootstrap support that would have to be incorrect. This is not uncommon (bootstrap support values only quantify sampling errors rather than accounting for anything more systematic, and can often be overly confident) and so the C. elegans & Drosophila genes have been classed as members of the orthogroup, but with low confidence.

**RefOG016.txt**

The RefOG tree is unrooted, it has been rooted on a Drosophila gene.

The newly inferred tree contains the target orthogroup and a number of further homologous genes. It has been rooted at the base of the clade containing the target orthogroup and a gene from the outgroup species Nematostella. It confirms the orthogroup from the original study with high bootstrap support.

**RefOG017.txt**

The RefOG tree is unrooted, it has been rooted at the base of the duplication separating the clade with the earliest diverging species (Ciona) from the other clade. This is the oldest branch in the tree.

The newly inferred tree contains an ancient duplication separating two sub-families with representatives from the earliest diverging species used in this study: Mnemiopsis leidyi & Trichoplax adhaerens. The tree shows the target orthogroup within its larger sub-family. The tree confirms the orthogroup identified in the original study and recovers genes from the outgroup species Mnemiopsis_leidyi, Trichoplax adhaerens, Nematostella vectensis. These demarcate the orthogroup. There is uncertainty over the gene Gallus_gallus_ENSGALP00000071704. They are all within the metazoan orthogroup but groups with 3 Mnemiopsis genes. However, its position is variable. In another tree it was shown as part of the sister bilaterian orthogroup. It has been excluded from this RefOG but with low certainty.

**RefOG018.txt**

RefOG tree is presented unrooted, it has been rooted on Danio & Tetraodon, the earliest divergence in the tree.

New tree with wider search shows clearly that the orthogroups goes back to the earlier diverging species: C. elegans & Drosophila. It has been rooted on the clade containing these two genes, the

Ciona gene is also in this clade, but such deviations from the expected topology are to be expected for some of these early branches, as the gene trees for other RefOGs in this study show.

**RefOG019.txt**

The RefOG tree is rooted on a Nematostella gene, there is very little resolution within the vertebrate clade due to lack of phylogenetic signal.

The newly inferred tree has genes from a number of outgroup species and from species diverging early within the orthogroup. They confirm the orthogroup from the original study, with some extra genes fitting clearly within the orthogroup (Pan & Danio, presumably new gene models) and missing a Ciona gene from the original study. This has been confirmed as a retired gene model (such a gene may exist, but it is no longer in the benchmarks input data and so shouldn't be an expected output).

**RefOG020.txt**

The RefOG has been rooted on a Nematostella gene. The tree suggests it may contain a duplication prior to the divergence of the Protostomes and Deuterostomes, in which case it would contain two orthogroups. Note, there are two copies for C. elegans and two vertebrate clades likely descended from this duplication.

The newly inferred tree (v1) contains the wider Metazoan orthogroup within its wider gene family. It has been rooted at the base of this Metazoan orthogroup. Using the tree to select the cut-off (rather than an e-value/bit score from hmmer) a new alignment and tree has been inferred on just the sequences from this Metazoan orthogroup. This tree (v2) shows that these two clades are separate bilaterian orthogroups with one clade containing representatives from the vertebrates C. elegans & Drosophila and the second clade containing genes from the vertebrates & C. elegans. As an aside, the wider tree actually appears to contain two Metazoan orthogroups and has been rooted as such, on the ancient duplication separating them.

The orthogroup containing Homo_sapiens_ENSP00000353654 has been arbitrarily taken as the target orthogroup.

**RefOG021.txt**

The RefOG tree is unrooted. It was rooted on the branch separating the Mus/Rattus genes from the remainder.

There are many homologous genes and there is no apparent evidence that only these 6 genes make up an orthogroup. The inferred tree shows at least three clades of genes that appear to have originated from duplications at the base of the mammals, well within the orthogroup.

What evidence does the original study give for the orthogroup being limited to just these genes?

- Category: Low quality MSA

- Occurs in Eukaryotes and Bacteria

- Protein involved in epidermis development

No information is given as to why this RefOG was so heavily circumscribed despite evidence to the contrary. A new tree has been inferred of 250 genes with hits to the hmm profile with e-values all better than 1.6e-9 from the larger gene family so as to identify the extent of this orthogroup within that gene family. The human gene ENSP00000357789 has been taken as the target since the tree appears to show that the gene Rattus_norvegicus_ENSRNOP00000053315 included in the original RefOG is only distantly related.

The orthogroups was identified as having a low quality MSA and so caution was exercised in interpreting the new tree. However, it was found to give a notably self-consistent picture of the gene family. Working back towards the root from the Homo & Pan genes there are a series of clades with the topology of each being consistent with the species tree and with generally high bootstrap support. This extends to the clade containing 125 genes all descended from duplications since the divergence of the vertebrates from the remaining species. Up to this point this is all with high certainty.

The next clade appears to mark the final extent of the orthogroup. It contains a gene from the outgroup species Nematostella vectensis, two genes from Drosophila and one from Branchiostoma as well as genes from the vertebrates. Thus, these genes appear to be a separate orthogroup. However, this is with low certainty. The branches lengths are long, there would appear to be moderate levels of gene duplication and loss and there is little structure to this subtree. It may be that these genes are not true homologs. The interpretation of these genes is unclear. They have been excluded from the orthogroup. The same is true of the remaining parts of the tree, suggesting that the tree does not require rooting elsewhere and that those sequences too may be false positives. With that said, the original 125 genes within the vertebrates are all part of the same orthogroup with high certainty.

**RefOG022.txt**

The RefOG tree is rooted on a Hydra magnipapillata gene although this is on a long branch and so the branch may not root the tree accurately.

The newly inferred tree shows three metazoan orthogroups in the gene family. It has been rooted at the base of the metazoan orthogroup for the target genes. This metazoan orthogroup is largely consistent with the original RefOG tree. As with the RefOG tree from the original study it contains earlier diverging genes within the clade identified as the orthogroup in the original study (Drosophila & Nematostella in this tree, Drosophila & C. elegans in the original RefOG tree).

If these genes are correctly placed then the clades contain two orthogroups, one (clade B) with only vertebrate genes in (the others lost post-duplication) and the other (clade A) with Deuterostomes, Protostomes & Cnidaria. Clade A has 100% bootstrap support, providing evidence that the duplication predated the orthogroup and that the clade B genes should be excluded. Clade A also has 100% bootstrap support in the tree from the original study. Arguing against this, the C. elegans gene is outside the well supported Clade A in the newly inferred tree whereas an earlier version of the gene model has it inside the well-supported clade A in the original RefOG tree (WBGene00001249.1 = W09C2.1a). This makes the status of the C. elegans gene uncertain, and potentially that of the Clade B genes.

A new tree (v2) has been inferred focusing on just the metazoan level orthogroup and rooted on Mnemiopsis_leidyi_ML409810a-PA & Trichoplax_adhaerens_TriadP56374 (the exact root is not clear, but doesn't affect the subsequent analysis). In this tree the C. elegans gene is part of the Clade B this time, despite 100% bootstrap support against this in the other two trees. Given that the bootstrap support values do not provide the confidence they would otherwise suggest, and given that the previous study identified clades A & B making up the orthogroup (which is at least a reasonable interpretation that matches the patterns of species presence in the appropriate sub-trees), both clades will be included in the orthogroup in this study. This is with low certainty.

**RefOG023.txt**

Roadmap of the Analysis performed on the RefOG

==============================================

Part 1: The gene sequence in most of the species is made up of a number of domains, and not all these domains have the same evolutionary history. The analysis of just this RefOG took well over a week. An alignment of the complete gene sequence gave a reasonable picture of how many of the genes within the vertebrates were related but the relationships between these larger clades of genes and between these larger clades and the outgroup species was unclear. This was because different parts had different histories and so concatenating them together produced inconsistent trees, as should be expected. Specifically, these clades moved relative to one another.

This calls into question whether the concept of orthogroup makes sense here. In the strict sense it doesn't, orthology should be traced for the each of the individual units that have a consistent history. Nevertheless, the final analysis did show that there was a core gene sequence that appeared to have a single evolutionary history and so (in the spirit of the original benchmarks) and orthogroup could be determined.

Part 2: Back to the analysis. The next step was to align the individual domains and examine the trees for these domains. There were 3 domains which were present in >90% of the genes likely constituting the orthogroup. Each of these domains had already been duplicated prior to the divergence of the Metazoa so each of these 3 trees had around 5 copies of the history of the gene tree, as shown by the subtrees reconstructed from the domain sequences, giving about 15 versions of the 'gene tree' with various levels of presence/absence and various levels of resolution. The sequences that these trees were determined from, however, were quite short and so although the general picture could be teased out for most of the genes in the vertebrates, the picture was hard to piece together nearer the base of the orthogroup.

Part 3: As hinted at by the analysis in part 2, there was a single evolutionary history that was shared by a number of the domains suggesting a core gene sequence inherited without other domains with different histories being inserted into it and appearing to be descended from the original core sequence. This was made up of the sequence of domains: VWD, C8, TIL, VWC, Pacifastin_I. A tree was inferred using this part of the gene sequence and allowed the orthogroup membership to be determined.

Part 1

======

(This can largely be ignored but is kept here for completeness)

The RefOG tree is rooted on a Hydra gene. A number of the branches have low support. It shows an orthogroup with a number of duplications around the vertebrates and mammals.

The newly inferred tree has been rooted on a well-defined metazoan-level orthogroup at some distance from the target genes. The tree recovers a Deuterostome clade with mostly high bootstrap support, as did the original RefOG tree. This is clade A (from ENSP00000382323 to ENSCINP00000014936). Earlier, there is a duplication giving rise to a second Deuterostome clade, clade B (from ENSCINP00000028688 to ENSP00000485659). Many of the genes in this clade were identified in the original study as members of the orthogroup, but strangely many have not, even though the tree gives very strong support for them being orthologs--they are in the same clade, the topology matches what would be expected from the species tree and the bootstrap support values are high.

The next earliest clade to diverge contains 4 genes from Nematostella & Mnemiopsis and so would appear to mark the extent of the orthogroup. This would require the gene to have been lost in the Protostomes, which is a possibility. The next clade is potentially and entire metazoan-level orthogroup, (clade C, from Trichoplax_adhaerens_TriadP21952 to Mus_musculus_ENSMUSP00000131401) which would support this. However, the branches here are short and could be hard to resolve in a large tree. Therefore, a second tree will be inferred on a clade encompassing all of the genes of interest, but smaller than the original tree. Original tree 458 genes, 89 genes so far identified as part of the orthogroup, the clade of 170 genes has been selected for the tree, going back to the clade of 8 Mnemiopsis genes.

The new tree (v2) has been rooted on these 8 Mnemiopsis genes. Unlike the first tree, it has placed two Nematostella genes as the sister clade to Clade B rather than to the combined Clade A+B. This has 80% bootstrap support, but the distribution of species makes this appear less likely (Clade B is vertebrates, Clade A is a vertebrate clade sister to Branchiostoma/Ciona clade). In the v1 tree, on the other hand, there was 83% bootstrap support for Clade A being the sister of Clade B with the Nematostella genes as the outgroup to these. The interpretation of the bilaterian-orthogroup suggested by the (v1) tree appears more parsimonious and better supported. It is also in closer agreement with the original study.

However, both trees agree on the 101 gene clade included the target orthogroup and stretching back to the first Mnemiopsis genes, the earliest diverging outgroup in the study. A such, one final tree will be inferred on just these genes and examined.

This new tree (v3) serves only to emphasise that the relationships of these clades are uncertain.

Part 2

======

(Informative towards the analysis performed in part 3, but can be skipped)

Trees were inferred for the domains VWD, C8 & TIL. These occur multiple times in each gene and from the gene trees this was the case in the ancestral gene sequence. Thus, these three trees give around 15 incomplete pictures of the evolution of the gene family between them. However, they didn't provide sufficient resolution near the base of the orthogroup.

Domain tree

-----------

Label with respect to v3 tree

C8 Part 1

---------

(((R2, P1),B3),B3),Mnemiopsis

Part 3:

(R12, B3), B3),P1,Mnemiopsis

VWD Part1

(R2, B)

VWD part 2:

(((B12,P2),B3),P1),R,

VWD Part 3:

(((R, Mnemiopsis), B124),B3)

Extending out from ENSP00000382323

---------------------------------

- Note, we should expect genes to be missing from some clades since that particular copy of the domain may have been lost

- Note, the numbers at the end of gene names are arbitrary labels added by me. They will differ between clades (the labels of different copies of the domain were assigned before the tree structure was known)

First confident clade:

Clade1_TIL.tree: Back to duplication prior to divergence of Danio

TIL 5 and TIL 4 both give the most complete picture and are also consistent with TIL 1, 2 & 3 for where they overlap.

TIL 5: unclear. There is a Mnemiopsis gene in the middle of the tree

TIL 4: Trichoplax gene in middle of tree.

TIL: ((blue,green),purple),red or (blue, red),green

C8 1: blue,(green, (red,purple)), black

Nematostella_vectensis_EDO44199 appears as an outgroup to

TIL 5 & TIL 3

and

C8 -3 (with mis-rooting)

C8 - 2.5 (blue clade is missing, it's well supported by all the other subtrees)

There seems to be good evidence for Blue, green/red/purple Drosophila, outgroup. With no outgroup coming between these in order to suggest a duplication prior to the orthogroup.

TIL 5: Matches this very well, other than Mnemiopsis_leidyi_ML231816a-PA_2 which falls in the middle of red. This never occurs elsewhere.

TIL 4: Same, but slightly disordered. Trichoplax_adhaerens_TriadP21952_4 in a different place in the middle.

There is an outgroup repeatedly coming in the middle of the red clade. But no other evidence, would expect other species to also come in the middle if this were the case.

VWD tree gives the clearest idea of the tree, but with certain clades missing.

VWD 4 ((blue,green):90,red),Drosophila

VWD 3 - poorly resolved

VWD 2 - poorly resolved

Part 3

======

(This section covers the final determination of the orthogroup membership). The corresponding files are 'v4'

A HMM was built around the sequence of pfam domains that were observed as common to a large number of the genes:

VWD

C8

TIL

VWC

Pacifastin_I

Examples of this domain were extracted from a localised clade of genes within the vertebrates since it was certain that they were all descended from an ancestral sequence within the orthogroup. Using the hits from the original search for all pfam domains, the sub-sequences within each gene corresponding to this sequence of domains was determined:

Homo_sapiens_ENSP00000382323:152-512

Homo_sapiens_ENSP00000447211:123-443

Danio_rerio_ENSDARP00000142662:133-494

Tetraodon_nigroviridis_ENSTNIP00000022420:39-360

The sequences were aligned with linsi, a HMM was built, it was searched against the genes and the parts of the genes that were hit were extracted. All target genes were hit by this hmm. These sequences were aligned using linsi and the tree was inferred using IQTREE on the untrimmed alignment.

Analysis of this tree

---------------------

Creating a combined profile for this sequence of 5 domains hit all 101 of the selected genes. A number of the genes scored multiple hits resulting in 498 hits in total. These hits were aligned, and the tree inferred. This tree traced the history of various copies of this domain within the gene family. Clades were identified in the unrooted tree which showed the history of homologous copies of the combined domain (see 5doms.linsi.fa.treefile.names.pdf). The clades were labelled clockwise from the top: A, B, C, D, E & F.

Clades B & D appeared to show the most complete history of the gene.

The tree was rooted at the bade of Clade B, and this clade was examined first.

Clade B

-------

The tree shows a number of duplications at the base of the vertebrates giving rise to 11 copies in Homo sapiens (plus a duplication of the domain within one of the genes giving rise to an extra copy of the domain in ENSP00000447211 & ENSP00000382323). This is made up of clades observed in all the previous single domain trees & these branches all have high bootstrap support. The bootstrap support at the base of this clade is 80%. Next there is a clade of 3 Branchiostoma genes (81% bootstrap support for this) and then a final clade of vertebrate genes containing the gene ENSP00000261405 (97% bootstrap supp port overall for this complete set of genes described so far). This is the (approximate) extent of the orthogroup. The next sister clades contain genes from Nematostella, Ciona, Branchiostoma, Mnemiopsis & Drosophila. There is then a further clade of vertebrate genes (containing Mus_musculus_ENSMUSP00000131401), followed by Trichoplax, followed by Mnemiopsis_leidyi_ML03227a-PA.

This is good evidence that the extent of the orthogroup has been (finally) located. Some further analysis is required to determine which of the genes from the 12 target species found around the base of the clade actually belong to the orthogroup and which diverged before the origin of the orthogroup. Before this, the next most promising clade will be examined to determine if it supports this overall analysis and, secondly, what additionally evidence it provides around the base of the orthogroup.

Clade D

-------

This clade has lower average bootstrap support and doesn't have any copies of its domains in humans (for example). It does not represent the core of gene sequence of the gene family.

Clade F (Mnemiopsis_leidyi_ML03226a-PA_4)

-------

This repeats the general picture of Clade B. The differences are: It shows that the duplication separating (ENSP00000382323, ENSP00000447211, ENSP00000261405) from (ENSP00000485659, ENSP00000490794, ENSP00000436812, ...., ENSP00000487059) actually occurred before the divergence of Branchiostoma rather than after (this was not visible from Clade B) with 92% bootstrap support.

Where does Drosophila fit into this gene family?

-----------------------------------------------

Clade A - This is made up of two metazoan-level orthogroups. A1: Drosophila_melanogaster_FBpp0075495_3 potentially duplicated (or this domain at least) and is in a separate bilaterian orthogroup supported by Branchiostoma genes in both apparent post-duplication clades (73%). A2 is less well resolved but potentially similar.

Clade B - it is present, one copy, likely part of the orthogroup rather than a duplication in the base of the Metazoa and then loss within the orthogroup.

Clade C - part of orthogroup but weak evidence as not outgroup species.

Clade D - similar to B? But evidence is not strong either way.

Clade E: Domain not present in Drosophila.

Clade F - this copy of the domain is not present

C. elegans & S. mansoni

-----------------------

Not present

So, the orthogroup contains a number of vertebrate clades that include 11 human genes. The gene was lost entirely in C. mansoni and C. elegans (likely two events if the species tree is correct) but not in Drosophila. However, there is a question as to whether the Drosophila gene is a descendant of a different ancestral gene from a duplication just before the ancestor of the bilateria or whether it is in the same orthogroup. In many ways, the concept of orthology and orthogroups is not appropriate for this gene, only for its individual domains. However, Clade B plus Clade F shows that there was a core gene sequence (made up of these two sequences) that was inherited, but was supplemented by additional copies of overlapping domains that had diverged further back in time. The orthogroup will be determined for this core sequence.

The genes for which membership needs to be resolved.

The clade (X): Monodelphis_domestica_ENSMODP00000003423
Tetraodon_nigroviridis_ENSTNIP00000011630 Canis_familiaris_ENSCAFP00000040196
Mus_musculus_ENSMUSP00000131401 Rattus_norvegicus_ENSRNOP00000037189
Danio_rerio_ENSDARP00000088544

The gene: Drosophila_melanogaster_FBpp0075495

Clade X there is 95% bootstrap support against it being in the same orthogroup in Clade F. 53% against in Clade A. Independent duplications in Clade D of no significance to the question. These genes will be excluded from the orthogroup.

Drosophila_melanogaster_FBpp0075495: Clade A in two places the evidence is against it being in the orthogroup. Clade B also 89% against. In other RefOGs the placement of an early diverging gene as diverging apparently before the outgroup species has not been taken as strong evidence for the gene's exclusion. This is because in many cases the tree shows an incorrect topology with no evidence for a duplication. The bootstrap only performs a resampling of the (limited) gene sequence data and so doesn't settle the question. However, in this case there is 1) good evidence for duplication events around the base of the orthogroup 2) Repeated placement of separate domains from the gene sequence outside the orthogroup. This provides evidence that the gene is not part of the orthogroup. This is with low confidence. The genes that have been ascertained to be part of the orthogroup are all with high confidence.

The members of the orthogroup were taken from Clade B & Clade F, they contained 71 & 73 genes respectively of which 70 were confirmed by both clades. Danio_rerio_ENSDARP00000150436_1 appears in the MSA to be in Clade B due to a duplication of a short part of Danio_rerio_ENSDARP00000130459_4 and its insertion into ENSDARP00000150436 and so it has been excluded. The three additional genes in Clade F (Gallus_gallus_ENSGALP00000051785, Pan_troglodytes_ENSPTRP00000092487 & Tetraodon_nigroviridis_ENSTNIP00000009471) all appear to be loss of one of the domains (the clade B) version but not the other (the Clade F) version, with the retention of the rest of the gene sequence and so are part of the orthogroup.

**RefOG024.txt**

The RefOG tree has two genes from outgroup species, it is rooted on one of these.

The inferred tree contains genes from the wider gene family. Its interpretation is not immediately apparent. The target orthogroup appears to be well-defined and bounded by a number of genes from the outgroup species. Nevertheless, other parts of the tree contain genes that don't fit well into expected clades and could potentially not be homologous. Although these genes may not be affecting the orthogroup of interest (which appears to be well resolved), a new tree with a more careful selection method for genes was used (the previous tree had a worst hit gene to the hmm profile of 0.0072).

A new tree was inferred a tree with 1.5x the number of genes as passed the e-value threshold as for the original RefOG tree so as to include the gene family context of the orthogroup. Thus:

47 genes in original tree, e-value of worst hit for profile was 3e-77. 69 genes from the new study had a hit better than or equal to this, selected the best 1.5x69 = 103 gene hits, with a worst hit to the hmm profile with a e-value of 1.8e-32 for being homologous.

The newly inferred tree contains a clade with a large expansion in gene copy number in C. elegans and Drosophila. These appear to be from outside the orthogroup. The tree has been rooted on the clade that contains the only three Mnemiopsis genes (a Ctenophore and the earliest diverging species in the tree) together with a single Drosophila gene. There is no clade containing only Mnemiopsis and no bipartition that appears to be a more ancient duplication, so this appears to be the best root for the tree. The Drosophila/C. elegans expansion appears to be from a separate orthogroup that doesn't retain any representatives from the Deuterostomes. The target orthogroup contains a clade of (outgroup) Nematostella genes diverging near its root and separating the orthogroup from this Drosophila/C. elegans expansion. There is also a pair of C. elegans & Drosophila genes which appear to be the representatives from the target orthogroup even though the topology does not match that expected for the species (these early branches are hard to resolve with single gene trees, as shown by many of the trees within this study). There is 98% bootstrap support for these genes belonging to the wider pre-bilateria orthogroup. This analysis supports the orthogroup inferred in the original study.

**RefOG025.txt**

The RefOG tree was rooted on a Nematostella vectensis gene.

The inferred tree contained the target orthogroup and a closely related clade which may be from the orthogroup or may be a closely related one, and then further orthogroup(s) separated by an ancient duplication from the target orthogroup. The presence of outgroup species bounding each of these clades confirmed that the duplication was ancient. The tree was rooted on the branch of this ancient duplication.

The inferred tree confirms the analysis from the original study in large part. The only gene in question is Ciona_intestinalis_ENSCINP00000003738. The inferred tree shows it as part of the second most closely related orthogroup to the target orthogroup, with these three orthogroups having diverged prior to the divergence of Trichoplax adhaerens from the remaining species.

The RefOG tree provides no evidence either way for this gene. If it were inside the orthogroup or outside the orthogroup both cases would produce the observed tree when rooted on the Nematostella as a result of the limited genes included in the tree since there are no earlier diverging genes which would allow these two possibilities to be distinguished between. The newly inferred tree does include such genes. The topology indicates that there was a duplication before the base of the orthogroup which gave rise to a Ciona gene and Branchiostoma lanceolatum gene. This would require a loss at the base of the Protostomes and another at the base of the vertebrates. Since there are already Ciona and Branchiostoma genes already in the orthogroup a duplication would similarly be required if these genes were part of the orthogroup followed a loss prior to the vertebrates. To place these genes in the orthogroup would involve contradicting a series of 4 bipartitions with high bootstrap support: 97%, 93% 100% and 100%. With this considered, these genes are unlikely to be part of the target orthogroup, but to have diverged earlier as show by the inferred tree.

**RefOG026.txt**

The RefOG tree is rooted on a Nematostella gene. It shows a single orthogroup with three duplication events at the base of the vertebrates.

The newly inferred tree has been rooted on an ancient gene duplication event prior to the origin of the target orthogroup. There are a number of updated or new gene models within the clade identified in the original study. It appears to confirm the orthogroup from the original study. The clade containing the two Drosophila and one C. elegans genes is shown as diverging prior to the divergence of the Cnidarian Nematostella. This bipartition has 89% bootstrap support. The two most likely possibilities are that these genes belong to the target bilaterian orthogroup or they belong to the sister orthogroup (containing

Homo_sapiens_ENSP00000347942). The previous study identified the genes as part of the target orthogroup. The 86% bootstrap support is in favour of this, although the topology of the tree suggests the opposite could be the case (with bootstrap support less than or equal to 10%). The gene have been assigned to the RefOG, as per the previous study but this is with low certainty.

**RefOG027.txt**

The RefOG tree is rooted on the Hydra magnipapillata gene. It appears to show a single orthogroup with a duplication in the vertebrates.

The newly inferred has tree contains a number of metazoan orthogroups within a larger gene family. It has been rooted at the base of the of the metazoan orthogroup for the target genes. It clearly confirms the orthogroup from the original study, with 100% bootstrap support at its base.

**RefOG028.txt**

The RefOG tree is rooted on a Hydra gene. The tree is confusing, it shows two clades separated with one of them containing a Drosophila gene. The tree makes sense as a single orthogroup if the

Drosophila gene is misplaced and actually diverged prior to the duplication. All the relevant branches that suggest otherwise have low bootstrap support.

The newly inferred tree has been rooted on a well-defined metazoan orthogroup that has 100% bootstrap support. The target orthogroup is within a metazoan clade with 98% bootstrap support and largely agrees with the RefOG from the original study. The Drosophila gene diverges prior to the duplication, which is more parsimonious. There are two genes with uncertain membership since they are shown diverging before the outgroup species: Tetraodon_nigroviridis_ENSTNIP00000006920 & Caenorhabditis_elegans_WBGene00002783.1. The orthogroup already includes the Tetraodon genes which would be expected in each of the clades within the orthogroup. The tree gives 94% bootstrap support for these genes not being members of the orthogroup. The have been excluded, but with low certainty.

**RefOG029.txt**

RefOG is rooted on the Cnidaria Nematostella vectensis. The tree looks clear, although the highest bipartition has bootstrap support of 1%, this is probably not an issue provided all the necessary genes were included. The tree has no branch lengths.

The inferred tree (v1) agrees with the RefOG tree in the broad outline. It splits the Ciona and Drosophila genes out and places them amongst the outgroups species but the clearest interpretation of these is that they belong in the orthogroup, as the RefOG also proposes.

In the inferred tree there is an extra clade of Homo, Pan, Mus & Rattus genes. These appear to be incorrect; they are a clade of a separate gene and shoehorning them into the tree has clearly disturbed the topology. Compare with the RefOG tree. The alignment confirms that this is the case.

A new tree (v2) has been inferred for these selected genes, it has been rooted on the Mnemiopsis gene.

The RefOG tree is correct here with moderate certainty.

**RefOG030.txt**

The RefOG tree has three genes from Nematostella/Hydra on which it is rooted.

The newly inferred tree has a number of outgroup genes delineating it, stretching back to Mnemiopsis leidyi. It has been rooted at the base of this metazoan orthogroup, on the duplication separating it from a homologous orthogroup. There is a Nematostella vectensis gene (EDO34323) in the outgroup part of the tree on a particularly long branch. It doesn't appear to have detrimentally affected the tree and the topology of the target clade, particularly around its root, is as expected. The tree clearly confirms the RefOG from the original study.

**RefOG031.txt**

The RefOG tree is rooted on Nematostella.

The newly inferred tree has been rooted at the base of one (or two) metazoan orthogroups containing the target bilaterian orthogroup. It shows the orthogroup from the original study to be correct. It has 99% bootstrap support and is demarcated by a number of outgroup genes. There are a number of extra genes corrected placed within the tree, likely from new annotations of the genomes.

**RefOG032.txt**

The RefOG tree is unrooted. It appears to show 3 bilaterian orthogroups that have been incorrectly identified as a single orthogroup. Each of these has a C. elegans gene, which is from the first diverging clade within the orthogroup thus showing the duplications occurred before the root of each orthogroup. It has been rooted on one of these orthogroups.

The newly inferred tree (v1) contains a number of metazoan orthogroups from the wider gene family. It has been rooted on a well-defined case which has outgroup genes from Trichoplax & Nematostella, and 100% bootstrap support. This is not to say that this is the actual root of the tree, but it does guarantee that the root is correctly positioned outside the target orthogroup and so the target orthogroup is correctly depicted in the rooted tree.

Interestingly, this tree does not have C. elegans genes delineating each of these three orthogroups, possibly because of updated gene models (but the genes appeared correct in the tree from the original study) or missed genes in this study. A manual search for missing C. elegans genes has been conducted and a tree focusing on just the clade of the genes in question plus close outgroup (to aid rooting) has been inferred. This outgroup clade was selected from the initial tree. The C. elegans gene in the tree from the original study and missing from the newly inferred tree is T14E8.1a. The updated ID for this gene is WBGene00020504. In fact, this gene is in the newly inferred tree but if shown, with 83% bootstrap support, to be a member of the bilaterian orthogroup containing Homo_sapiens_ENSP00000431885. I.e. the orthogroup selected to be the outgroup for the new tree. Returning to the initial inferred tree from this study, it presents good evidence that this C. elegans gene belongs to that orthogroup. I will return to the second tree soon.

Given this, the RefOG tree was re-rooted on the C. elegans gene, given the good evidence that it is an outgroup to all the remaining genes in the tree. This still shows there to be three bilaterian orthogroups of which two have representatives from the Protostomes and Deuterostomes and one only has Deuterostomes.

The second, more focused tree (v2) inferred for this study confirms that the RefOG from the original study is in fact 3 bilaterian orthogroups. The orthogroup containing Homo_sapiens_ENSP00000478721 has been chosen as the representative for the benchmark as it is the most clearly defined.

**RefOG033.txt**

The RefOG tree is unrooted, it appears to show a single orthogroup with 4 duplications mostly around the base of the vertebrates. It has been rooted on one of these vertebrate orthogroups.

The newly inferred tree (v0) shows the target orthogroup as part of the larger gene family, it has been rooted at the base of the metazoan orthogroup containing the target genes. Some of the clades in the tree look like they may be incomplete (as can be expected if only genes meeting a certain e-value threshold are used). This leaves uncertain the relationship to the target orthogroup of the 7 genes in the clade containing Homo_sapiens_ENSP00000216862. A new tree has been inferred with a wider net so as to better understand the position of these genes.

This tree (v1) contains approximately 4 metazoan orthogroups in a gene family that includes the target bilaterian orthogroup. It has been rooted on the most distant of these from the target orthogroup, which is also the most clearly defined. The target orthogroup has two outgroup species genes and has very high bootstrap support throughout including 100% at its base. It clearly confirms the orthogroup identified in the original study and shows the clade containing Homo_sapiens_ENSP00000216862 is from a separate orthogroup.

**RefOG034.txt**

The RefOG tree has been rooted on Ciona.

The newly inferred tree has been rooted at the base of a metazoan orthogroup at a duplication with 98% bootstrap support for the bipartition. It has 4 outgroup species genes. It confirms the orthogroup from the original study.

**RefOG035.txt**

The RefOG tree is rooted on Nematostella, it has no branch lengths.

The newly inferred tree covers numerous orthogroups within the gene family. The target orthogroup has a number of outgroup genes from earlier diverging Metazoa, it has been rooted on a bipartitions with 99% support representing a duplication which separates this and 2 further metazoan orthogroups from the rest of the gene family. The tree shows outgroup genes for both the target orthogroup and the adjacent orthogroup (containing Homo_sapiens_ENSP00000481105), confirming the delineation of the orthogroup from the original study. There is a Ciona gene outside of this clade in the tree for this orthogroup. The evidence that it does not belong in the target orthogroup is that there is a bipartition with 60% bootstrap support separating it from the orthogroup and the tree also shows that there have been many duplications within this family near the base of the Metazoa, so its position within the tree is not unreasonable. It could also potentially belong to one of the neighbouring clades.

**RefOG036.txt**

The RefOG tree is rooted on Nematostella. It shows a single orthogroup with a number of internal duplications at the base of the vertebrates.

The inferred tree shows the target orthogroup with non-bilaterian outgroup genes and the gibbering orthogroup that diverged near the base of the Metazoa. It confirms the orthogroup membership from the original study. There is are two outgroup genes between the Ciona gene and the rest of the clade, but the support values are low and so, as with the original study, this gene has been included in the orthogroup.

**RefOG037.txt**

The RefOG tree is unrooted. It shows two clades separated by a duplication either before the bilaterian common ancestor (and hence actually made up of two orthogroups) or after the divergence of the Protostomes from the Deuterostomes (and hence a single orthogroup). It contains a single Nematostella gene, on which the tree has now been rooted. This clearly shows the two clades.

The newly inferred tree (v1) extends much further across the gene family so as to give a clearer picture of the target orthogroup in its larger context. It shows at least three metazoan-level orthogroups. It has been rooted at the base of the metazoan orthogroup containing the target bilaterian-level orthogroup. This clade contains two genes from the Ctenophore Mnemiopsis leidyi (the earliest diverging species included) as well as representatives from Nematostella and Trichoplax. The tree shows that the RefOG from the previous study may be a single bilaterian-level orthogroup or may be two. The tree has been used to select a more focused set of (68) genes including these one or two bilaterian orthogroups plus the outgroup species.

This second tree (v2) has been rooted on the two Mnemiopsis leidyi genes. The presence of 4 genes from some early-branching but in-group species in a clade with Trichoplax adhaerens & Nematostella vectensis does not affect the following interpretation of the target orthogroup. They are most likely non-separable from the true outgroup due to the difficulty of resolving these earliest branches. This complication is avoided by the tree from the previous study only due to the use of a single gene. The newly inferred tree has on average higher bootstrap support as well as better species sampling.

Whether the two clades belong to the same or different bilaterian orthogroups depends on whether the duplication occurred before or after the divergence of the Protostomes from the Deuterostomes. One of the clades has both Protostomes from the Deuterostomes as well as the earlier diverging Nematostella. This suggests these are two orthogroups, unless the tree inference is in error and these genes in fact diverged before the duplication. This possibility has to be considered since there are only Deuterostomes in the second clade. The first clade has 88% bootstrap support. This is strong but not overwhelming evidence, but the topology is the only evidence there is and it is favour of these clades being separate orthogroups. The RefOG has thus been identified as only the clade containing the gene Drosophila_melanogaster_FBpp0112980. This is more likely correct than incorrect, but with low confidence.

**RefOG038.txt**

The RefOG tree has been rooted on Nematostella. It has no branch lengths.

The newly inferred tree covers a number of metazoan orthogroups. Each of the closest orthogroups to the target orthogroup are clearly demarcated by their own outgroup genes. The tree has been rooted on a duplication separating these orthogroups from a number of orthogroups included the target.

The clade containing the target genes stretches appears to have a single gene from the Protostomes (Schistosoma_mansoni_Smp_179370) and many genes from the Deuterostomes. If this is the case then this is the complete orthogroup and it is in agreement with the original study. This clade has 100% bootstrap support.

The two closely related orthogroups are each bounded by their own outgroups and suggest a duplication at the base of the Metazoa, confirming the target orthogroup identified in the original study.

**RefOG039.txt**

RefOG tree is unrooted. Rooted on C. elegans.

The newly inferred tree contains the larger gene family of which the orthogroup is a part. Rooted at the base of the metazoan orthogroup containing the target. It has outgroup genes from Nematostella vectensis & Mnemiopsis leidyi. There is some doubt over whether the clade containing Homo_sapiens_ENSP00000418754 is a duplication inside the orthogroup. There are no Protostomes in both clades (C. elegans in one and S. mansoni in the other) however the tree topology shows that the duplication predated the orthogroup and the bootstrap support values are high: 98% & 100% and so the orthogroup has been identified as the same clade as the original RefOG. This is with moderate confidence.

**RefOG040.txt**

The RefOG tree has been rooted on the Drosophila gene.

The inferred tree recovers a number of metazoan orthogroups. It has been rooted on the duplication branch separating the target orthogroup and its (presumably) two closest sister orthogroups from the remaining orthogroups. It agrees with the original RefOG orthogroup with numerous outgroup species demarcating the clade.

**RefOG041.txt**

The RefOG tree is rooted on Hydra_magnipapillata_Hma2.224352.

The newly inferred tree was rooted on the earliest diverging outgroup gene, Mnemiopsis_leidyi_ML00976a-PA. It confirmed the orthogroup from the original study. There are few AA substitutions within the mammalian clade, emphasising the length of the branches for Canis_familiaris_ENSCAFP00000031345, Rattus_norvegicus_ENSRNOP00000005511, Pan_troglodytes_ENSPTRP00000091393 but the actual branch length for this clade is not long and the hmm profile search showed these genes to be homologous with high certainty & similar to the other genes in the orthogroup.

**RefOG042.txt**

The RefOG tree has been rooted on a Nematostella gene.

The newly inferred tree (v1) contained two genes from species that had been added to help with the analysis, but were likely false positive homologs, Trichoplax_adhaerens_TriadP63087 & Branchiostoma_lanceolatum_BL23244_evm4 which both had e-values for the hmm profile > 1e-4.

Examination of the MSA provided further evidence for this. They were removed from the MSA and the tree was re-inferred.

The second newly inferred tree (v2) was rooted on the ancient duplication separating the two clades. The topology of these two clades suggested that the comparatively long branch connecting them had intercepted each of the sub-trees deep within these sub-trees rather than at their respective roots. This can occur with a long branch intercepting a sub-tree with many short branches, it is equivalent to the inaccuracy possible when rooting a tree with a too-distant outgroup.

The two clades are clearly distinct orthogroups, each having multiple representatives from the outgroup species. This confirms the focusing of attention on just one of these as the target clade in the original study. To confirm the orthogroup membership within this clade a third tree was inferred with just the genes from this clade.

The third newly inferred tree (v3) showed the clade to be composed of two bilaterian orthogroups, duplicating within the Metazoa and both having outgroup genes from the Cnidaria. The new RefOG was arbitrarily chosen to be the one containing Homo_sapiens_ENSP00000347324.

**RefOG043.txt**

The RefOG tree is rooted on an outgroup gene, there is another within the clade, but I agree with the interpretation that this is misplaced.

The inferred tree agrees with the RefOG, with the updating of gene models. It has been rooted at the base of the Metazoan orthogroup.

**RefOG044.txt**

The RefOG is rooted on a Hydra magnipapillata gene.

The newly inferred tree has extra outgroup genes from Nematostella, Trichoplax & Mnemiopsis. They show that the RefOG from the previous study incorrectly combined genes from two bilateria orthogroups. This reiterates the danger of rooting a tree on a single outgroup gene when there is a duplication adjacent to the root since two different cases can't be distinguished: (Outgroup,(Clade1, Clade2)); and ((Clade1, Outgroup), Clade2);

The newly inferred tree clearly shows two orthogroups with outgroups for both with 95% bootstrap support for them being separate bilaterian orthogroups. The one containing Homo_sapiens_ENSP00000349577 has been taken to be the target. The one containing Homo_sapiens_ENSP00000301175 has been labelled RefOG044b.

**RefOG045.txt**

The RefOG tree is rooted on Nematostella.

The newly inferred tree (v1) shows a large gene family made up of many bilaterian orthogroups. It is largely in agreement on the membership of the target orthogroup. There are two genes, in Gallus gallus and Danio rerio which are most likely members of another orthogroup, but it would be safe to infer a tree more closely focused on the target orthogroup in order to be sure.

This more focused tree (v2) confirms that these two genes are not members of the orthogroup.

**RefOG046.txt**

The RefOG is rooted on Hydra.

The bipartitions in the RefOG and the inferred tree have mostly low support. The newly inferred tree has been rooted on Mnemiopsis leidyi.

The orthogroup is easily identified and the RefOG from the previous study is in good agreement with the inferred tree (with updating of gene models).

**RefOG047.txt**

The RefOG tree is rooted on a Hydra magnipapillata gene.

The newly inferred tree has been rooted on the Mnemiopsis leidyi gene. The extra species used in this tree have not all been arranged as would be expected from the species tree topology (Nematostella & Trichoplax), but the bootstrap support is not high for these placements either--they don't give evidence from the C. elegans genes being excluded from the orthogroup. Overall, it shows that there is a single orthogroup, in agreement with the original RefOG tree.

**RefOG048.txt**

The RefOG tree is unrooted, it has been rooted on the Nematostella vectensis gene.

The inferred tree was poorly resolved, with a number of polytomies within the orthogroup. Inspection of the RefOG tree showed this also be the case for that tree. These are largely due to lack of changes between sequences within the orthogroup whereas there does appear to be sufficient resolution closer to the boundary of the orthogroup. The newly inferred tree has been rooted on Mnemiopsis leidyi with the next diverging species being Nematostella & Trichoplax, indicating good resolution here.

There appear to be a large number of genes from within the orthogroup not identified in the previous study which should be added to the RefOG with high certainty. The limit of the bilaterian orthogroup are easy to distinguish and coincides with the limit identified in the previous study.

**RefOG049.txt**

The RefOG tree is unrooted. It has been rooted on Danio & Tetraodon.

The inferred tree clearly shows this orthogroup and at least some of the members of the next most closely related orthogroup. Both orthogroups have a clear outgroup gene from the Ctenophore Mnemiopsis leidyi with high bootstrap support. This gives high confidence in the orthogroup membership. The tree has been rooted on the branch separating these two orthogroups. The tree is in good agreement with the orthogroup from the previous study.

**RefOG050.txt**

The RefOG is rooted on Nematostella.

The inferred tree has been rooted on the Mnemiopsis leidyi gene and is in good agreement with the original RefOG.

**RefOG051.txt**

The RefOG tree is rooted on a Nematostella gene.

The inferred tree contains three orthogroups, it has been rooted on a duplication separating the most clearly demarcated of these from the other two. The tree largely supports the RefOG orthogroup. There is a gene in the original RefOG from a separate clade that had not been fully recovered in the previous study (Tetraodon_nigroviridis_ENSTNIP00000019325). The question is whether the duplication giving rise to this clade was before or after the origin of the target orthogroup. The tree shows it as before with moderate bootstrap support, 85%. And the topology and pattern of species presence/absence are both consistent with a duplication prior to the divergence of Branchiostoma so this clade has been included in the orthogroup. It is probable that Caenorhabditis_elegans_WBGene00020557.1 also belongs to the orthogroup. There is 77% bootstrap support for it being in the same clade and there is not a copy of a C. elegans gene in the clade with Schistosoma & Drosophila despite no other putative gene loss events. With moderate certainty the gene belongs in the clade with those genes and has been included in the orthogroup.

**RefOG052.txt**

The RefOG has been rooted on the branch separating the mammals from Danio/Tetraodon.

The inferred tree shows that this clade reaches back further, to the starlet sea anemone, Nematostella vectensis. It has been rooted on this clade. One of the sister orthogroups is also demarcated by the outgroup species Trichoplax adhaerens. The tree shows, with 100% bootstrap support, that the orthogroup contains an extra pair of genes from Danio & Tetraodon and a gene from Ciona.

**RefOG053.txt**

The RefOG is rooted on the Nematostella vectensis (Cnidaria) gene.

The orthogroup is well demarcated in the tree (v1) by genes from the outgroup species Nematostella vectensis & Trichoplax adhaerens. It has been rooted at the base of this larger metazoan-orthogroup. The inferred tree supports the orthogroup membership from the original study with updated gene models.

However, genes could have potentially been incorrectly excluded by the e-value threshold for inclusion so a new tree (v2) with all hits better than or equal to 0.01 were included. No trimming was used on the MSA. This tree, with 46 genes rather than 24, presents a more complex picture. As before the main clade back to Ciona intestinalis is well-supported. Also, as for the v1 tree the evidence supports Drosophila_melanogaster_FBpp0297348 also being a member of the bilaterian-level orthogroup. The genes that are less clear are Gallus_gallus_ENSGALP00000073380 and the three C. elegans genes.

The v1 tree provides a clear picture with the two Trichoplax genes as the outgroup & two Nematostella genes diverging next, as expected. The C. elegans genes are in a clade with the Schistosoma mansoni genes, as expected and EDO37889 appears to just be slightly displaced in the tree.

Looking at the unrooted v2 tree, it two clear clades. One involves the target orthogroup and also stretches back to the Nematostella and Trichoplax genes. The other is distant & includes ENSP00000367459 with similar species representation. Between these two clades are a number of genes on long branches and in no biologically expected tree topology. Are these spurious, false positive homologs or are the part of the gene family? And correctly placed in the tree? The position of the C. elegans genes in relation to the Nematostella & Trichoplax genes suggests they are not part of the orthogroup, with at least 70% bootstrap support. This is also in agreement with the v1 tree once it is rooted as per the v2 tree (which has more information as to the root). Conversely, Gallus_gallus_ENSGALP00000073380 appears to be part of the orthogroup. Both of these decisions are with only moderate confidence.

**RefOG054.txt**

The RefOG tree is unrooted. It has been rooted on Ciona.

Most bipartitions have high support but there is no outgroup.

The newly inferred tree has a number of outgroup genes from Nematostella vectensis & Mnemiopsis leidyi providing a highly supported outgroup clade. The tree has been rooted on the Mnemiopsis gene. The tree shows clearly that there was a duplication within the Deuterostomes giving rise to the additional clade within the orthogroup, 96% bootstrap support (containing ENSP00000392762). The orthogroup also clearly extends to the Protostomes, with the topology closely matching what would be expected and with 96% bootstrap support.

**RefOG055.txt**

The RefOG tree is unrooted, it has been rooted on the Ciona genes.

The inferred tree has been rooted on a group of Nematostella genes (v1). It is in good agreement with the RefOG from the previous study. The inferred tree has two genes inserted within this orthogroup on long branches: Canis_familiaris_ENSCAFP00000051664 & Ciona_intestinalis_ENSCINP00000012544. The topology doesn't correspond to what would be expected for these genes nor their point of insertion within the tree. It is unlikely that these are correctly placed within the tree. The MSA supports the theory that these are false positives. And the hmmer profile search gave an e-value of 5.7e-63 for a closest neighbour within the tree versus 0.0052 & 0.00021 for these genes. These genes will not be included in the orthogroup, in agreement with the original study. In further support for this, a previous tree included Mus_musculus_ENSMUSP00000041113 in a clade with these two genes. It has a similar match to the hmm profile. In this tree, that gene is now in a different location in the tree, outside the orthogroup. These two likely false positive homologs should not have been included in the tree. A new tree has now been inferred without these two genes (v2). A lighter trimming was also used since the available sequence is particularly short.

**RefOG056.txt**

The RefOG is unrooted. Rooted on Danio & Tetraodon (there is a Drosophila gene in the tree, but it is on a long branch in the mammalian clade).

The inferred tree supports the RefOG with relatively high certainty. Mnemiopsis leidyi & Branchiostoma lanceolatum genes were identified and each have 100% bootstrap support for their respective clades. The only concern is that there are Drosophila & Tetraodon genes diverging from the target orthogroup clade before Mnemiopsis leidyi. On reflection, these genes probably belong to this orthogroup, but this is with low certainty. The clade containing the Mnemiopsis gene only has 51% bootstrap support, it is more likely that the Drosophila & Tetraodon belong in this orthogroup.

The only gene of doubt is Danio_rerio_ENSDARP00000104764. It is in the RefOG tree and fits perfectly but did not get found with the hmmer search for inclusion in the newly inferred tree. Note that the RefOG contained 10 genes whereas the top 55 hmmer hits have been used for the newly inferred tree so this is surprising.

Ensembl says this gene has been retired. The tree suggests it is ok, but if it's not in the input dataset then it should not be included in the expected benchmark output from these datasets (regardless of whether it is actually a real gene or not).

RefOG is correct with updating of gene models.

**RefOG057.txt**

RefOG tree is rooted on the Nematostella vectensis. The tree looks good and the important branches for delineating the orthogroup have high bootstrap support.

The inferred tree supports the delineation of the original RefOG. A few genes appear to have been missed from the original RefOG (from Danio, Mus) but these may be new gene models. There was an extra Ciona gene in the RefOG but this has been retired.

**RefOG058.txt**

The RefOG tree is unrooted, it has been rooted on the branch separating the mammals from Danio/Tetraodon.

An initial tree aiming for approximately triple the number of genes in the tree from the original study (see methods) did not give a complete picture of the orthogroup and so a new, larger tree was inferred with a more relaxed e-value cut-off (v1) with 250 genes. The inferred gene tree shows that this gene family is complex. It has been rooted on an ancient duplication distant from the target orthogroup and near the root of the Metazoa to give a clear view of the target orthogroup and what appears to be its most closely related sister orthogroup. The newly inferred tree gives strong evidence for an additional clade (ENSP00000367013) of genes within this orthogroup originating from a duplication at the base of the vertebrates with 99% bootstrap support. There is also a second duplication giving rise to a further clade in the vertebrates (ENSP00000349324). The genes Branchiostoma_lanceolatum_BL08616_cuf2 & Nematostella_vectensis_EDO33270 are most likely misplaced, they have poorer hits to the hmm profile than genes in either of the two surrounding clades.

However, this is very hard to have any certainty over. Another possibility is that there are genes closely related to BL08616 & EDO33270 which have been excluded from the tree, resulting in only a partial picture. A new tree (v2) with an HMMER e-value cut-off of 0.001 has been inferred, this is highly unlikely to result in any false positive homologs. As can sometimes occur, this has resulted in a rearrangement of some of the clades. Both v1 & v2 trees need to be considered....

To recap, the RefOG from the original study had 9 genes centred around ENSP00000162749. With very high confidence this is incorrect--there was a duplication within the orthogroup giving an additional clade of 8 genes around ENSP00000367013 also in the RefOG. The question that is unresolved is whether the other gene duplication in the gene family occurred within the target orthogroup or before it. In general, it is hard to make sense of either of the trees in terms of the distribution of the various species in the tree, particularly the outgroup species. What could be ascertained with reasonable confidence from the unrooted v2 tree was the target orthogroup of interest could be rooted at the base of a clade of 93 genes. A new tree (v3) was inferred from these genes. This tree has the advantages that 1) it has, with reasonable confidence, all the genes in it that need to be considered since the clade has been determined from tree inference rather than using HMMER e-values 2) It is no larger than necessary, and so the genes of interest are more likely to be correctly resolved than either of the larger trees, additionally, the MSA is likely to be of higher quality.

The v3 tree can be securely rooted on a group of 3 Mnemiopsis genes (the fourth Mnemiopsis gene is also close to this root). This tree (at last!) is clear: the arrangement of the species throughout the tree is as expected from the species tree and, in particular, this is apparent when there are gene duplication events. In these cases, the distribution of species either side of the gene duplication event is consistent as is the distribution of species on the branches prior to the gene duplication event.

This tree shows, working backwards in time: The original clade A (ENSP00000162749), the clade B (ENSP00000367013) that resulted from a gene duplication event prior to the divergence of the vertebrates, another gene duplication event giving rise to the clade C (Mus_musculus_ENSMUSP00000119790), one further gene duplication event still in the ancestor of the vertebrates giving rise to the clade D of 30 genes (ENSP00000498466 amongst other human genes), the speciation event giving the genes from the outgroup species Nematostella vectensis (EDO37507 & EDO33270) and marking the boundary of the bilaterian-level target orthogroup.

This final branch has 43% bootstrap support, suggesting that some of genes which have been excluded/included could actually be members/non-members of the orthogroup. However, the remainder of the clades in the tree are all consistent with this current interpretation, indicating a gene duplication event prior to the orthogroup (and prior to the divergence of Nematostella) giving rise to the sister clade of 33 genes, which itself contains another gene duplication event prior to the divergence of Nematostella to result in two other bilaterian level orthogroups (around ENSP00000172229 & ENSP00000266557 respectively). Furthermore, the delineation of the orthogroup is a very large improvement on that achieved in the previous study which included just 9 genes in the vertebrates, with no evidence to support marking those genes off as a complete orthogroup.

**RefOG059.txt**

The RefOG tree is rooted on a Nematostella gene.

The inferred tree has been rooted on a Mnemiopsis leidyi gene and is in good agreement with the RefOG tree.

**RefOG060.txt**

The RefOG tree is rooted on a Hydra (fresh-water polyp) gene. It does not have branch lengths.

The method for selecting genes for the initial tree produced a tree (v1) with 425 genes of which the target orthogroup was only a small part. A new tree with the best 250 hits was inferred (v2). The inferred tree shows a clear orthogroup which can be rooted on the Drosophila/C. elegans clade. The RefOG from the previous study has also included a vertebrate clade containing Mus_musculus_ENSMUSP00000023400, but this is clearly a part of a separate orthogroup that also includes representatives from C. elegans & Drosophila as well as all sampled species from the Deuterostomes. The true orthogroup is well supported by the inferred tree, with the topology matching well the known species topology and with 100% bootstrap support for the bipartition at its root.

The intention of the v2 tree was that the tree inference would have been better able to explore the part of tree space relevant to the RefOG more easily without the extra ~200 genes from outside the RefOG also included. However, the tighter e-value threshold excluded Caenorhabditis_elegans_WBGene00017400.1 from consideration. The v1 tree shows 100% bootstrap support for this being a paralog of Caenorhabditis_elegans_WBGene00007670.1, and a member of the orthogroup. This highlights the importance of a lenient e-value threshold if the orthogroup is to be determined by the gene tree rather than which genes reach an arbitrary e-value threshold.

**RefOG061.txt**

The RefOG tree is unrooted, it has been rooted on the most ancient branch, separating the mammals from Danio/Tetraodon.

The inferred tree has been rooted at the base of the clade of genes. It shows the two clades from the original study (A containing ENSP00000369004 & B ENSP00000346634) but additionally three Danio genes clearly orthologs of this first gene. Also an additional clade, C, containing ENSP00000435210 more closely related to the B than B is to A, with 100% bootstrap support.

There is some uncertainty as the rooting of the tree is not secure. This is because homology with the remaining genes in the tree is far from certain and so point the branch from those trees inserts into the tree for the target orthogroup is not necessarily the true root. With this unreliable rooting accounted for, the position of the three clades A, B & C as duplications within the orthogroup still remains the best interpretation of the tree.

**RefOG062.txt**

The RefOG has been rooted on an outgroup gene from Monosiga brevicollis.

The inferred tree contains genes from a number of orthogroups, it has been rooted on one of the ancient duplications separating these orthogroups. The inferred tree largely supports the RefOG from the previous study. There are a number of genes outside the main clade (88% bootstrap support) and these have very low sequence similarity scores (hmmer) compared to the genes which are clearly in the orthogroup so it is likely then are not members of this orthogroup.

**RefOG063.txt**

The RefOG tree is unrooted. It has been rooted on the Tetraodon gene.

Searching for genes for the tree revealed some uncertainty over which genes should and should not be included in the gene tree. In most RefOGs an e-value of 0.001 has been used. All members of the orthogroups have comfortably obtained a hit to the hmmer profile at least this good. For this orthogroup this doesn't appear to be the case.

The gene sequences are very short and the range of e-values is continuous suggesting that there could be many true positive hits at high e-values. On the other hand, adding non-homologous sequences to the tree will force the tree to place these non-homologous genes somewhere (they may be detectable as misplaced) and could affect the position of other genes (this is more troubling and harder to detect).

The number of hits achieved at different e-values:

54 for 0.001 - v1

58 for 0.01

67 for 0.1 - v2

107 for 0.99

From the trees there are 19 genes clearly within the orthogroup. This increases to 23 when including a clade (ENSP00000220812) with an uncertain relationship to the orthogroup (tree v1). The membership of the main clade of 23 genes does not change as the e-value cut-off is changed from 1e-3 to 0.99. The second clade (ENSP00000220812) expands to give a reliable, separate clade of genes when an e-value of 0.1 is used (v2). It is unclear what the relationship is between the two clades--this needs to be established. The tree also shows many duplicates in Branchiostoma lanceolatum, an in-group species that diverged early within the in-group clade added to the analysis to aid the investigating of the orthogroups. It has been rooted on this clade of genes. This is the earliest diverging species in the family other than a Nematostella vectensis gene. This species is an outgroup and is nested within the Branchiostoma clade of genes. Rooting on this gene instead would not affect the interpretation. Trees v1 & v2 have been rooted on the branch separating the Branchiostoma clade from the remaining genes. This is not to suggest this is the root, which has not been established, all that is asserted at this point is that the root lies outside of the main clade of 23 genes, which is sufficient for the analysis.

Because of uncertainty over which genes to include in the tree, a new HMM profile was inferred based only around those genes which are clearly members of the orthogroup. The scores/e-values for the hits will then give an indication of how closely the genes are to this clade. The 19 genes in the two clades around ENSP00000295619 and ENSP00000271331 were aligned using mafft linsi and a hmm profile built. This was searched against the proteomes and genes achieving an e-value of 0.1 or better were included in the tree. 52 genes hit this profile with e>=0.001 and 87 with e>=0.1 (remember, the sequences are about 110 AA long so e-values will be low. The e<=0.001 tree was missing a number of crucial genes (for example, the clade around Homo_sapiens_ENSP00000220812 was incomplete) and so the e<=0.1 was used. This is tree v3.

The rooting of this tree will affect the interpretation. It has two outgroup genes and two Drosophila genes. For the rooting, the tree can be viewed, unrooted, in three parts. Part A contains the two vertebrate clades already identified as clearly members of the target orthogroup (around ENSP00000271331 & ENSP00000295619). Part B contains a large number of Branchiostoma genes & a gene from Mnemiopsis. Part C contains a set of vertebrate genes containing 3 human genes. The root can clearly be excluded from A. It's unlikely to be in B for two reasons.

Firstly, the gene tree makes a lot more sense with the root at the base of C as it then shows an ancient duplication, and then in one half the divergence of the Branchiostoma clade B from the vertebrate clade A while in the other half Drosophila diverges first then there are a series of Deuterostome clades. It topology is not perfect in this case: Drosophila_melanogaster_FBpp0072926 is in the Branchiostoma clade and Mnemiopsis_leidyi_ML070257a-PA appears in a location within the putative target orthogroup (most likely, its position should be shifted a short distance in the unrooted tree to lie just inside the C clade rather than just inside the B clade). If the root were within the B clade there is no logical interpretation of the tree. At best, the root would be on the Mnemiopsis gene and the Branchiostoma clade would diverge first and then there would be a gene duplication event giving rise to clades with a Drosophila gene at the base of one of them.

Secondly, the hmmer search suggested that the Branchiostoma clade of genes is a lot more closely related to Clade A than any of the Clade C genes are to Clade A. As such, the rooting on Clade C has been taken as the most probable.

The rooted tree shows the two vertebrate clades of A as clear members of the orthogroup, as before. The Branchiostoma genes are also part of the orthogroup, although these are only here to help resolve the trees, Branchiostoma is not one of the 12 target species. The gene Mnemiopsis_leidyi_ML070257a-PA should likely be the other side of the bipartition that has 41% bootstrap support. The only genes that are uncertain are Tetraodon_nigroviridis_ENSTNIP00000006119 & Drosophila_melanogaster_FBpp0072926 which are both shown by the tree as members of the orthogroup, but with poor support. The gene Drosophila_melanogaster_FBpp0309033 is also of interest, it is in a logical place within the tree for NOT being part of the orthogroup but by a small distance and only with 41% bootstrap support separating it from a logical position within the orthogroup. The simplest explanation is in agreement with the tree--Tetraodon_nigroviridis_ENSTNIP00000006119 & Drosophila_melanogaster_FBpp0072926 are in the orthogroup and the other Drosophila gene is part of the sister orthogroup. ENSTNIP00000006119 would have arisen from a gene duplication event and then been lost twice, once in Danio rerio and once before the divergence of Gallus gallus. All the trees up to this point have consistently placed it in this location.

**RefOG064.txt**

The RefOG tree is unrooted. It has been rooted on C. elegans & Drosophila.

The inferred tree identified additional outgroup genes and agrees with the RefOG orthogroup.

The gene Caenorhabditis_elegans_WBGene00010139.1 appears amongst the orthogroup, but out of place in the inferred tree. A duplication could have occurred before the origin of the orthogroup. There are also two genes from Trichoplax, weakly supporting a duplication before the orthogroup. It has been included in this orthogroup, but with low certainty.

**RefOG065.txt**

The data from the original study does not include a tree from RefOG065, the tree labelled as such is a duplicate of the tree for RefOG002. Nevertheless, the HMM profile is correct.

The newly inferred tree (v1) contains the target orthogroup together with outgroup species and shows two or more gene duplication events prior to the MRCA of the Metazoa. It has been rooted at the base of the metazoan-level orthogroup containing the target bilaterian orthogroup. It shows a clade of almost entirely single-copy genes with representatives from all but 2 of the 12 species. The presence of the 3 outgroup species genes together in the tree but not at the root of the metazoan-level orthogroup suggests the long branch from the distant metazoan orthogroup may not have intersected the short branches at the root of the orthogroup of interest. A new tree was inferred on just the metazoan-level orthogroup containing the target bilaterian orthogroup.

This second tree (v2) has been rooted on the Mnemiopsis gene, the first diverging species and reproduces the expected topology, confirming the hypothesis for the initial tree. The orthogroup is fairly straightforward, the only question with respect to genes from the 12 species of interest is Drosophila_melanogaster_FBpp0081863. A literal reading of the tree shows with (only) 49% bootstrap support that it diverged prior to the origin of the orthogroup. There has been a gene duplication event near the root since there are two Drosophila genes. There are also two Branchiostoma genes. The best interpretation is that there was a duplication prior to the origin of the orthogroup, since the multiple copies are seen in both the Protostomes and Deuterostomes, and then losses in each of these clades. Thus Branchiostoma_lanceolatum_BL00270_evm0 and Drosophila_melanogaster_FBpp0081863 are from a separate orthogroup. This more closely matches the topology of the tree than two separate gene duplication events (and losses) within the orthogroup. This is with only with low confidence.

**RefOG066.txt**

The RefOG tree was unrooted. It has been rooted on the Nematostella gene.

The inferred tree supports the RefOG tree. A Trichoplax and Nematostella gene are show within the clade, suggesting they mark the extent of the orthogroup. There isn't evidence for these genes being the result of duplication and selective loss, whereas many gene trees fail to correctly resolve the relationships of genes from these species--the exact topology of the species tree around the base of the Metazoa continues to be further refined by studies with larger species and gene sampling. The Trichoplax gene is concerning since it has high bootstrap support, but the topology of the tree above this strongly supports this gene being an incorrect insertion into an otherwise single-copy clade of genes rather than a duplication. For this reason, it is relatively likely that these genes do belong to the orthogroup, as proposed by the original study.

**RefOG067.txt**

The RefOG tree is rooted on a Nematostella gene.

The inferred tree agrees on the orthogroup, some genes are misplaced in the tree but not more than would be expected for these deeper branches and given the bootstrap values/length of the MSA.

**RefOG068.txt**

RefOG is rooted on a Nematostella vectensis gene. The identification of the RefOG in the original study is incorrect, details as to the probable cause are given at the end as they are not relevant to the identification of the actual orthogroup but are included for completeness.

The inferred tree clearly shows two metazoan-level orthogroups, with each containing genes from the outgroup species Mnemiopsis leidyi, Schistosoma mansoni & Trichoplax adhaerens. It is not entirely clear whether a small number of genes belong in the orthogroup or not: Drosophila_melanogaster_FBpp0309618, Ciona_intestinalis_ENSCINP00000027090, Ciona_intestinalis_ENSCINP00000001707, Caenorhabditis_elegans_B0511 & Danio_rerio_ENSDARP00000131597. They all appear in the tree to have diverged from the remaining orthogroup genes before the divergence of the outgroup genes Mnemiopsis_leidyi_ML040024a-PA & Nematostella_vectensis_EDO36444.

1. Are some/all of these genes incorrectly included in the tree and do not belong in the orthogroup?

2. Remnants of a duplication event predating the orthogroup and with the gene lost in the remaining species?

3. Members of the orthogroup that have been forced out of the clade in the tree as a result of artefacts: atypical sequence divergence, poor alignment, truncated gene models etc?

With low certainty, all of these genes other than Danio_rerio_ENSDARP00000131597 belong in the orthogroup. The tree excludes this gene with high bootstrap support. Hard to resolve short branches has most likely resulted in the remaining four genes being placed amongst the outgroup genes to a greater or lesser extent.

More notes:

There are three closely related vertebrate clades, all with high bootstrap support. The two Branchiostoma lanceolatum diverge prior to these clades, which is the correct location given the species tree.

From the MSA EDO36444 & EDO36443 appear to be two halves of the same gene and Mnemiopsis_leidyi_ML040024a-PA appears to be a faulty gene model and on a long branch. It is only included to serve as an outgroup and there are already a number of other closely related outgroup genes at this point so I have removed this in case it is interfering with the position of the gene it has been placed as a sister to.

The tree, after these corrections to the alignment, shows more clearly that the 2 Ciona and the Drosophila gene belong in the orthogroup. The C. elegans gene is more questionable, but it has a very truncated sequence and so its placement is not so surprising.

Why were genes from two orthogroups incorrectly identified as belonging to a single orthogroup?

The inferred tree shows that there are two separate orthogroups that have been labelled as a single orthogroup in the RefOG. Inclusion of more genes from outgroup species would have shown this. The tree has been rooted on a single gene from Nematostella despite the fact that a branch on the path between it and e.g. Homo_sapiens_ENSP00000385713 (most likely the branch directly above the Nematostella gene) is a duplication event more ancient than the divergence of the Nematostella gene from the clade containing Homo_sapiens_ENSP00000347931. Note, the topology of the inferred unrooted tree "(Ingroup1, Outgroup, Ingroup2);" is insufficient to distinguish between these two cases, although viewing the tree as a 'Radial Phylogram' in Dendroscope would give a strong indication of what had gone wrong here.

**RefOG069.txt**

The RefOG tree is unrooted. It has been rooted on the duplication separating the two vertebrate clades. It only contains representatives from the vertebrates, there is no apparent evidence that the orthogroup only extends back to this duplication at the base of the vertebrates and not further.

The newly inferred tree shows an additional duplication at the base of the vertebrates and a further clade of genes in this orthogroup. This tree only contains genes which obtained a HMMER hit better than an e-value of 0.001. Given that the Pan troglodytes gene ENSPTRP00000050875 only achieves and e-value of 0.00089 but clearly sits within this third clade, there is good reason to try a tree with more (also likely) homologous sequences to see where they appear in the tree. The e-value for the two Pan genes from the original study were 3.4e-46 and 6.3e-41, so that does suggest caution should be exercised when considering the genes from the third clade as also part of the orthogroup. It is necessary to weight up the two possibilities:

1. Was there a duplication within the orthogroup and the sequences are short/divergent and so diverged considerably in the 3rd clade. It would be useful to search with a single sequence as a seed rather than a hmm profile to gauge how quickly sequence similarity falls off within this orthogroup. With a hmm profile built on the sequences themselves (as is the case for the two original Pan genes), the e-values will appear misleadingly strong.

2. The homology suggested by hmmer is misleading. Or is it very distant and there was a matched pattern of three losses (Protostomes, Branchiostoma, Ciona) in all three clades of genes.

Query ENSMUSP00000061185:

| | | | | | | | | | | | |
|---|---|---|---|---|---|---|---|---|---|---|---|
| ENSMUSP00000061185 | ENSMUSP00000061185 | 100.0 | 187 | 0 | 0 | 1 | 187 | 1 | 187 | 1.0e-84 | 310.1 |
| ENSMUSP00000061185 | ENSRNOP00000016953 | 87.2 | 187 | 24 | 0 | 1 | 187 | 1 | 187 | 2.1e-79 | 292.4 |
| ENSMUSP00000061185 | ENSPTRP00000034753 | 76.5 | 196 | 37 | 1 | 1 | 187 | 1 | 196 | 1.1e-62 | 236.9 |
| ENSMUSP00000061185 | ENSP00000276571 | 76.5 | 196 | 37 | 1 | 1 | 187 | 1 | 196 | 1.2e-62 | 236.9 |
| ENSMUSP00000061185 | ENSCAFP00000040748 | 71.4 | 196 | 47 | 1 | 1 | 187 | 1 | 196 | 6.3e-60 | 227.6 |
| ENSMUSP00000061185 | ENSMODP00000036360 | 64.4 | 194 | 62 | 3 | 1 | 187 | 1 | 194 | 3.0e-46 | 182.2 |
| ENSMUSP00000061185 | ENSGALP00000050250 | 55.9 | 188 | 61 | 5 | 1 | 187 | 1 | 167 | 1.2e-32 | 136.7 |
| ENSMUSP00000061185 | ENSDARP00000114719 | 46.3 | 188 | 74 | 4 | 1 | 187 | 1 | 162 | 2.2e-26 | 116.7 |
| ENSMUSP00000061185 | ENSTNIP00000000470 | 40.0 | 190 | 84 | 5 | 1 | 187 | 1 | 163 | 1.9e-23 | 106.3 |
| ENSMUSP00000061185 | ENSDARP00000118835 | 41.2 | 136 | 44 | 4 | 54 | 187 | 54 | 155 | 2.6e-14 | 76.6 |

### Approx boundary of first clade

| | | | | | | | | | | | |
|---|---|---|---|---|---|---|---|---|---|---|---|
| ENSMUSP00000061185 | ENSGALP00000073211 | 62.0 | 50 | 19 | 0 | 138 | 187 | 58 | 107 | 5.5e-09 | 58.2 |
| ENSMUSP00000061185 | ENSDARP00000019621 | 54.2 | 48 | 22 | 0 | 140 | 187 | 91 | 138 | 1.6e-08 | 57.4 |
| ENSMUSP00000061185 | ENSRNOP00000008037 | 48.1 | 54 | 27 | 1 | 134 | 187 | 70 | 122 | 1.6e-07 | 53.5 |
| ENSMUSP00000061185 | ENSMUSP00000035321 | 48.1 | 54 | 27 | 1 | 134 | 187 | 70 | 122 | 1.8e-07 | 53.5 |
| ENSMUSP00000061185 | ENSCAFP00000031248 | 46.3 | 67 | 35 | 1 | 121 | 187 | 61 | 126 | 1.6e-07 | 53.5 |
| ENSMUSP00000061185 | ENSMODP00000028973 | 45.5 | 44 | 24 | 0 | 144 | 187 | 87 | 130 | 8.9e-06 | 47.8 |
| ENSMUSP00000061185 | ENSPTRP00000020205 | 44.8 | 67 | 33 | 2 | 124 | 187 | 59 | 124 | 2.1e-05 | 46.6 |
| ENSMUSP00000061185 | ENSP00000296099 | 44.8 | 67 | 33 | 2 | 124 | 187 | 59 | 124 | 2.3e-05 | 46.6 |

ENSMUSP00000061185 ENSGALP00000071657 44.7 38 21 0 150 187 63 100 5.3e-04 41.6

### Approx boundary of second clade

This strongly suggests that the detectable level of homology falls off quickly for this gene family and that the e-values for the third clade of genes when searching with a hmm profile are not any lower than would be expected for genes in the same orthogroup separated by a duplication prior the divergence of the vertebrates but within the orthogroup. All the genes in the newly inferred tree are members of the orthogroup. I.e. it is larger than the originally proposed RefOG69.

**RefOG070.txt**

The RefOG tree is rooted on the Nematostella vectensis gene XP_001631125.1 there is also a Hydra magnipapillata gene XP_002162003.1. There are two further genes from Hydra and Nematostella shown in the tree as sister to the Drosophila gene. The tree has no branch lengths.

The inferred tree shows clearly that this is indeed the orthogroup. There was a duplication prior to divergence of the Metazoa giving rise to a second, sister orthogroup. Both these orthogroups are clearly demarcated by representatives from the outgroups Trichoplax adhaerens, Nematostella vectensis, Mnemiopsis leidyi & Schistosoma mansoni around the base of each orthogroup.
